# Different functional connectivity gradients reflect aging and Alzheimer’s disease

**DOI:** 10.1101/2025.05.22.655469

**Authors:** Jonathan Rittmo, Nicolai Franzmeier, Olof Strandberg, Léa Chauveau, Theodore D. Satterthwaite, Laura EM Wisse, Nicola Spotorno, Harry H Behjat, Amir Dehsarvi, Danielle van Westen, Toomas Erik Anijärv, the Alzheimer’s Disease Neuroimaging Initiative, Sebastian Palmqvist, Shorena Janelidze, Erik Stomrud, Rik Ossenkoppele, Niklas Mattsson-Carlgren, Oskar Hansson, Jacob W Vogel

## Abstract

Aging and Alzheimer’s disease (AD) are accompanied by alterations to large-scale communication patterns in the brain, which can be tracked in vivo using functional connectivity (FC). The location, direction and relevance of these changes remain widely debated, though they are rarely studied in the context of whole-cortex communication dynamics. In two independent cohorts (BioFINDER-2, N=973; ADNI, N=129), we show that FC changes associated with aging and AD are strongly aligned with separate fundamental axes of hierarchical brain communication. Early accumulation of AD pathology and subsequent cognitive decline are both linked to functional change along the sensory-association axis. Meanwhile, age-related functional changes occur along the representation-executive axis consistently throughout the adult lifespan. These findings together suggest AD and aging both alter major but orthogonal functional pathways in the brain. More broadly, our findings position whole-brain connectivity dynamics as a unifying framework for interpreting functional changes across the adult lifespan.

## Main

Aging and Alzheimer’s disease (AD) are both associated with changes in functional brain networks – groups of regions that typically communicate with one another and support cognition. However, the precise nature of these changes – their spatiotemporal dynamics, and whether they reflect compensatory adaptations, pathological disruptions, or both – remains a topic of active debate^1,2^. Answering these questions is not only important for understanding disease biology, but could enable tailored clinical strategies that address age-independent mechanisms of AD. AD is the leading cause of dementia and is characterized by pathological accumulation of amyloid-β and tau proteins. These proteins spread through the brain in characteristic patterns^3,4^, leading to neurodegeneration and cognitive decline. Functional connectivity (FC), the temporal synchrony of activity between brain regions, may be a mechanism facilitating the spread of pathology, a system affected by pathology, or an interplay of both^5^. However, normal or healthy aging is also associated with altered functional connectivity^6–8^. These age and AD-related alterations have been proposed both as mechanisms to sustain cognitive function as the brain is challenged by stressors, and as manifestations of pathological network breakdown^2,9–12^.

FC changes in aging and AD have commonly been studied through two approaches. One approach focuses on regional connectivity changes, such as increases or decreases within or between specific regions or networks, or between regions of interest and the rest of the brain^13,14^. Although some consistent effects have been reported, such as reduced FC within the default mode network in healthy aging^7^, the overall picture is more complex. Studies of aging, especially in the context of AD, frequently report mixed patterns of hyper- and hypoconnectivity^15–19^. Such mixed findings may arise from several sources. Contributing factors include the predominantly cross-sectional nature of existing studies, unmeasured pathology, region- or network-specific analytical approaches that may miss broader spatial patterns, and small sample sizes that may not represent the broader population. An additional possibility is that the relationship between connectivity and age or disease progression is nonlinear – that is, hyper-may shift to hypoconnectivity (or vice versa) depending on age or disease severity. While initial hyperconnectivity is often attributed to adaptive cognitive compensation^2,12^, the relationship between connectivity and cognition remains unclear. The common regional approach of FC research allows for precise and granular analyses, but is inherently limited in providing a more holistic perspective on functional changes in healthy and (AD) pathological aging.

The other common approach for studying FC changes in aging and AD adopts a more holistic view, analyzing connectivity patterns by decomposing connectivity matrices into components of variance often referred to as ‘gradients’^20^. These components organize brain regions along axes of functional similarity, representing whole-cortex patterns of brain organization. The primary components capture the largest variations in the data, arranging regions along a continuum that groups areas with similar brain activity together, while placing regions with the greatest dissimilarities at opposite ends. The sensory–association (SA) axis^21^, which is often the primary variance component of FC^20,22^, groups unimodal sensory and motor cortices (such as occipital, precentral, and postcentral regions) on one end, and spans toward transmodal association areas supporting higher-order cognition on the other (including the medial prefrontal cortex [mPFC], posterior cingulate and precuneus, angular and inferior parietal cortices, and lateral and anterior temporal regions). Typically, the second component spans visual to motor cortices (VM axis), while the third component tends to separate different transmodal systems. Specifically, the executive lateral fronto-parietal system lies on one end and parts of the default mode network, including mPFC and anterior temporal regions, lies on the other^23^. This latter axis has been referred to as a “representation-modulation” axis^24^, or simply a “multiple-demand” gradient^25,26^. Based on this prior literature and our own characterization (see Figure E1) we will refer to it as a “representational-executive” (RE) axis.

**Figure E1:**
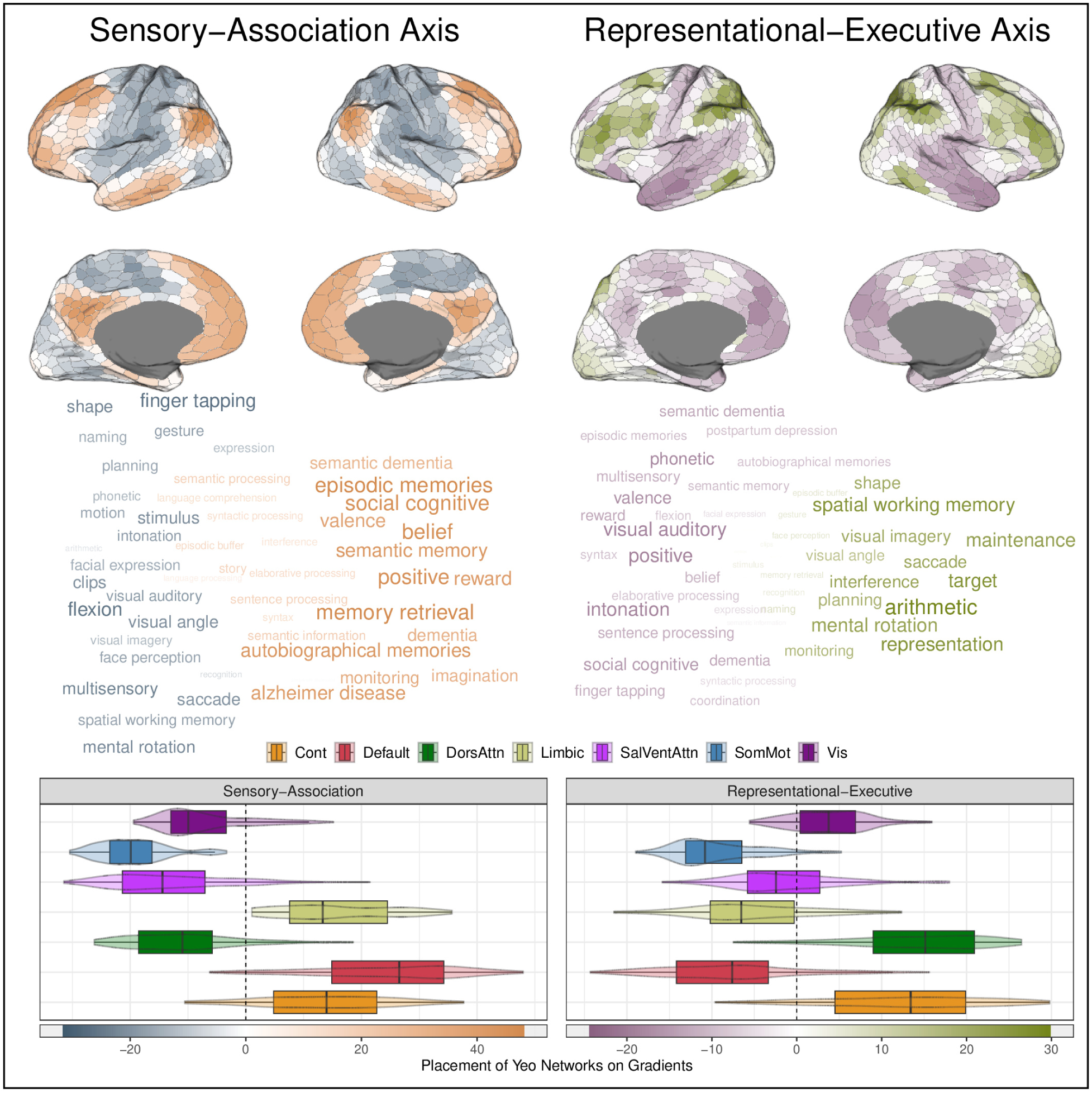
Functional connectivity gradients capture spatial organization and cognitive specialization across the cortex. Top: Spatial distribution of gradient scores for the SA axis (left) and the RE axis (right) respectively, derived from biofinder. Middle: Meta-analytic decoding was performed using the NeuroQuery database^35^ to identify cognitive terms associated with each Schaefer-1000 parcel. For each parcel, regional associations with NeuroQuery terms were estimated using the ROIAssociationDecoder in NiMARE, and results were aggregated across parcels. The resulting term associations were filtered to include only cognitive concepts listed in the Cognitive Atlas^36^. Words were assigned to a gradient side (e.g. sensory or association depending on the gradient of interest) decided by their average gradient value, and within each side the 25 most frequently mentioned words (i.e., the highest number of parcel associations) were retained. For each term, we counted the number of parcels it was associated with and its mean gradient value, assigned it to a gradient side, and retained the 25 most frequently associated terms per side. The terms are visualised along an x-axis of, and weighted both in size and colour by, their mean gradient score. Bottom: Distribution of gradient scores across the 7 Yeo networks.

A main feature of the gradients lies in capturing the brain’s functional architecture as continuous, overlapping maps ranking regions by their degree of involvement in distinct functional domains. But most prior work has studied gradients in terms of how their expression changes, for example becoming more pronounced during development^27,28^ and less pronounced with aging or Alzheimer’s disease^29,30^. In contrast, we do not treat functional gradients as the primary outcome of interest. Instead, we use normative functional gradients as a reference framework to interpret how connectivity changes in aging and AD are spatially organized. Recent work has demonstrated that neurodegenerative variation can be embedded within low-dimensional manifolds closely aligned with the canonical functional gradients. For example, FDG-PET derived axes have been shown to somewhat differentially capture aging- and AD related variance,^31^ and large-scale connectivity alterations in neurodegeneration have been shown to have implicit gradient structure^32^. Building on these insights, we ask whether functional connectivity itself reorganizes along normative gradients across the AD continuum, and whether such alignment differs between aging and AD-related processes.

To do so, we quantify region-wise connectivity using nodal affinity. This measure quantifies the similarity of each region’s functional connectivity profile to all others. Variation in affinity reflects if the region is more or less distinct in its communication pattern. In this way we integrate the two dominant approaches outlined above, combining regional granularity with a whole-cortex functional perspective. However, capturing a comprehensive picture of how FCS is altered by aging and AD requires consideration of not only where these changes occur, but also when. To address this, we model nonlinear region-wise trajectories of connectivity across age and along the AD continuum using a continuous biomarker-based measure of pathology, then examine how longitudinal increases in pathology affect concurrent changes in connectivity. Finally, to determine whether these connectivity changes relate to cognition independently of age and pathology, we conduct separate analyses in cognitively unimpaired and cognitively impaired individuals. Altogether, this study provides a framework that leverages group-level functional gradients and a continuous biomarker-based pathology measure to examine the spatial distribution, cognitive relevance, and temporal progression of FC changes in aging and AD.

## Results

We analyzed resting-state fMRI data from the BioFINDER-2 cohort (973 participants) with complete data (CSF Aβ42/40, tau PET and fMRI scans; see Table 1 for participant characteristics). After parcellating the cortex into 1000 regions^33^, FC was quantified using nodal affinity, the parcel-wise average of connectivity similarity to all other parcels. Henceforth, we focus on this functional connectivity similarity (FCS) as our central FC measure of interest, see Figure 1 A and Methods. Nodal affinity was used as the outcome in parcel-wise regression models (see Methods), with the t-values of the terms of interest projected onto the cortical surface, resulting in “t-maps” (Figure 1 A, C). Our goal was to assess the degree to which these t-maps aligned with the principal axes (i.e. gradients) of functional organization (Figure 1 C). The gradients of interest were defined as the first three principal components of the averaged connectomes from healthy controls (Figure 1 B; see Figure E10 for comparisons of different methods to derive gradients, and explained variance of the derived components). Supplementary analyses (Table S1) with gradients derived using several methods (including the traditional diffusion map embedding from^20^) confirmed that this methodological choice did not substantially impact the results. Due to a non-existent or modest and non-reproducible relationship between the t-maps and Gradient 2 (see Figure S1), this gradient was left out of subsequent analyses for brevity. Details about the anatomical, functional and network coverage of the SA and RE axis are presented in Figure E1. The age- and AD-related t-maps were spatially compared to the SA and RE axis, using Pearson correlation to quantify their relationship with statistical significance assessed using a spin test (one-sided, uncorrected; see Methods) to account for spatial autocorrelation^34^. We interpret gradient alignment primarily based on effect-size magnitude and consistency across cohorts and analyses; modest but statistically significant correlations are reported but not emphasized.

**Figure 1:**
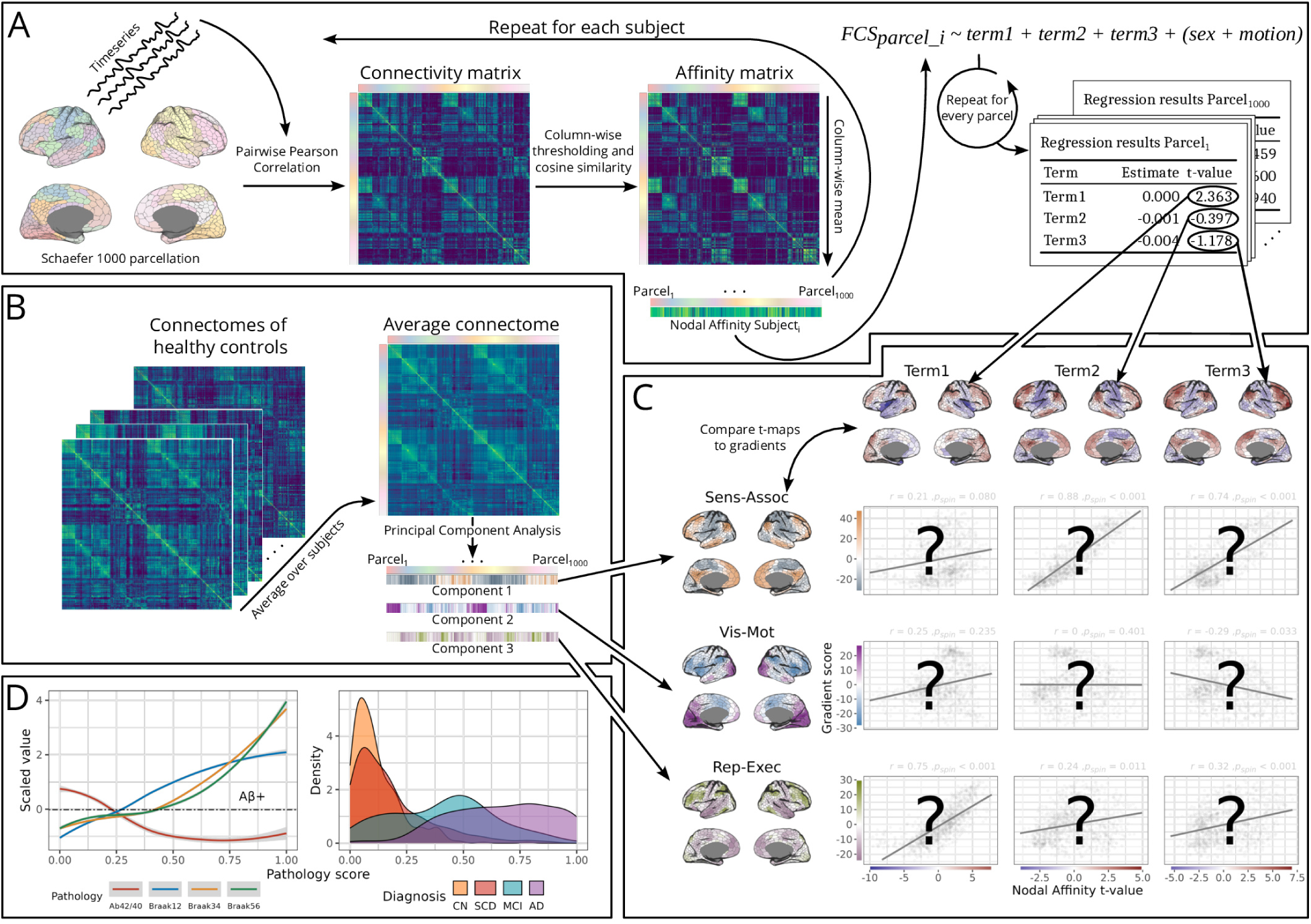
Mapping changes of functional connectivity similarity to cortical gradients. (A) Resting-state fMRI data for each subject were parcellated into 1000 regions^33^. Timeseries from these parcels underwent pairwise Pearson correlation. Cosine pairwise similarity between the connectivity profiles of the parcels were then calculated and averaged for each parcel, resulting in one FCS value per parcel and subject^20^. This served as the outcome in parcel-wise regression models, with t-values of the terms of interest (e.g., age, diagnosis) mapped onto the cortical surface in (C), forming t-maps. (B) Gradients were derived by averaging connectivity matrices of healthy controls and calculating principal components. The first three components/gradients were retained but only the first (SA axis) and third (RE axis) further analysed. (C) T-maps are compared to gradients and their relationship quantified using spin tests (one-sided; see Methods). (D) Participants were given a pathology score by mapping them nonlinearly onto a continuous pathology trajectory using Aβ and tau biomarkers using the SCORPIUS method^37^, providing a gradated and biological alternative to clinical diagnoses for subsequent analyses (see Methods for details). The relationship (LOESS curves) between the pathology score and the scaled biomarkers used to derive it is shown to the left, and distributions of clinical diagnosis groups to the right.

**Table 1:**
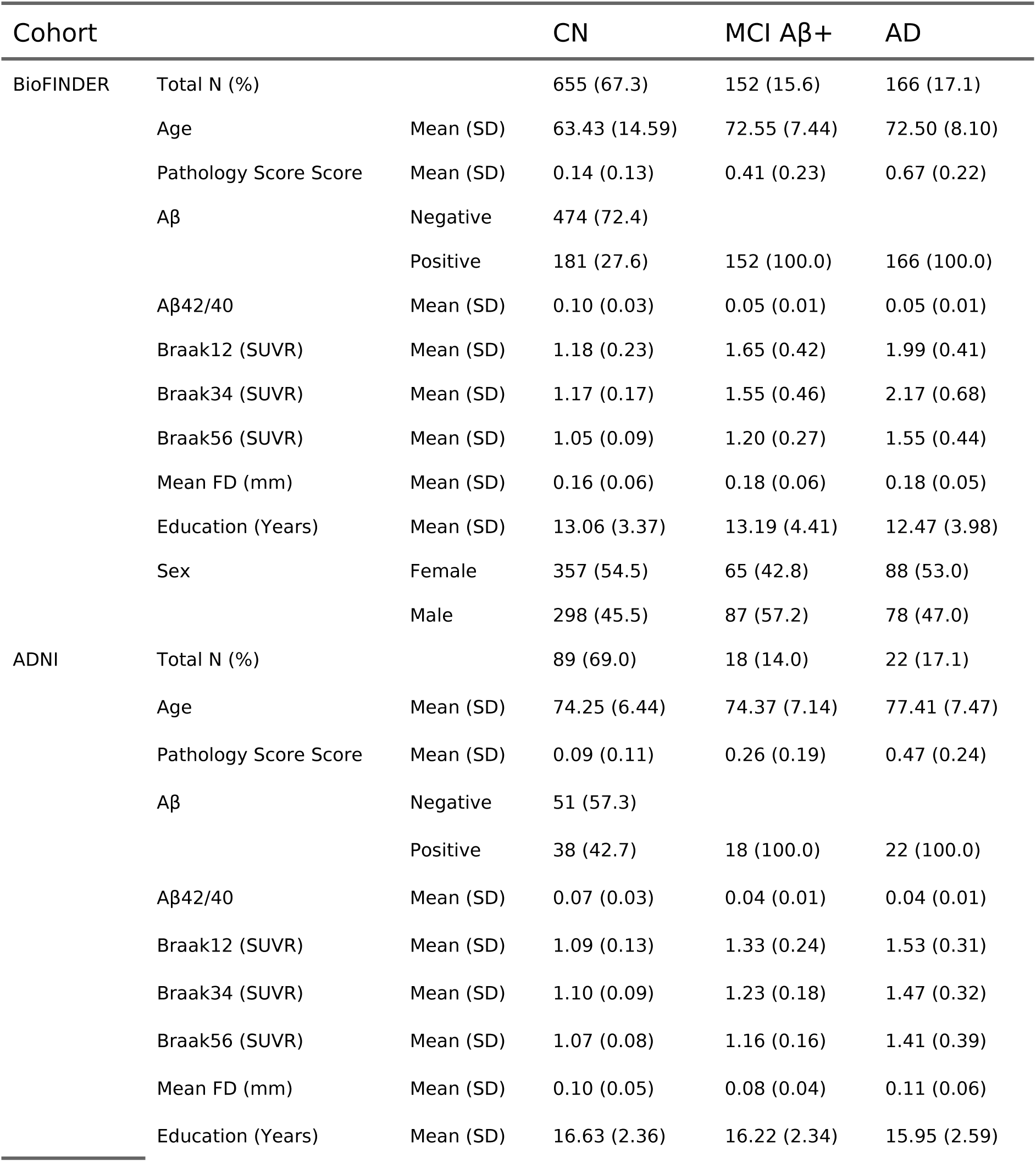
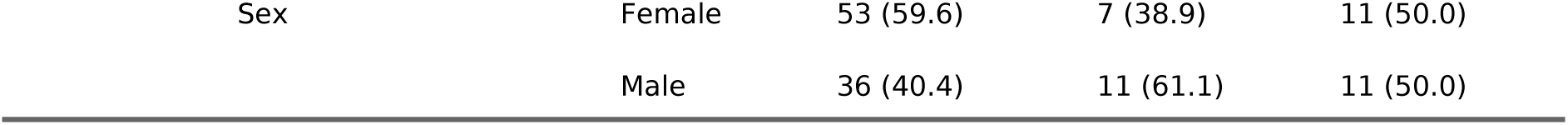
Cross-sectional characteristics in BioFINDER and ADNI. Pathology scores were calculated by non-linearly mapping individuals’s CSF Aβ42/40 ratio and tau PET SUVR from Braak I-II, III-IV, V-VI regions onto a trajectory from 0 to 1 where 0 indicates no pathology. Abbreviations: FD = frame displacement (scanner motion); SD = standard deviation, CN = cognitively normal; MCI = mild cognitive impairment; AD = Alzheimer’s disease.

In studies comparing aging and AD, it is common to rely on clinical diagnostic criteria. However, these criteria may confound pathological processes with cognitive factors (e.g., resilience). To isolate pathological AD effects, we used a continuous measure of pathology severity by nonlinearly mapping participants onto a continuous trajectory based on CSF Aβ42/40 ratio and tau PET using the SCORPIUS method^37^. This continuous pathology measure was used instead of clinical diagnoses to quantify AD disease progression in all subsequent analyses (but see Figure E2 for an analysis with clinical diagnoses). In Figure 1 D, the relationship between this composite pathology score and its constituents is visualized, along with its distribution over clinical groups.

**Figure E2:**
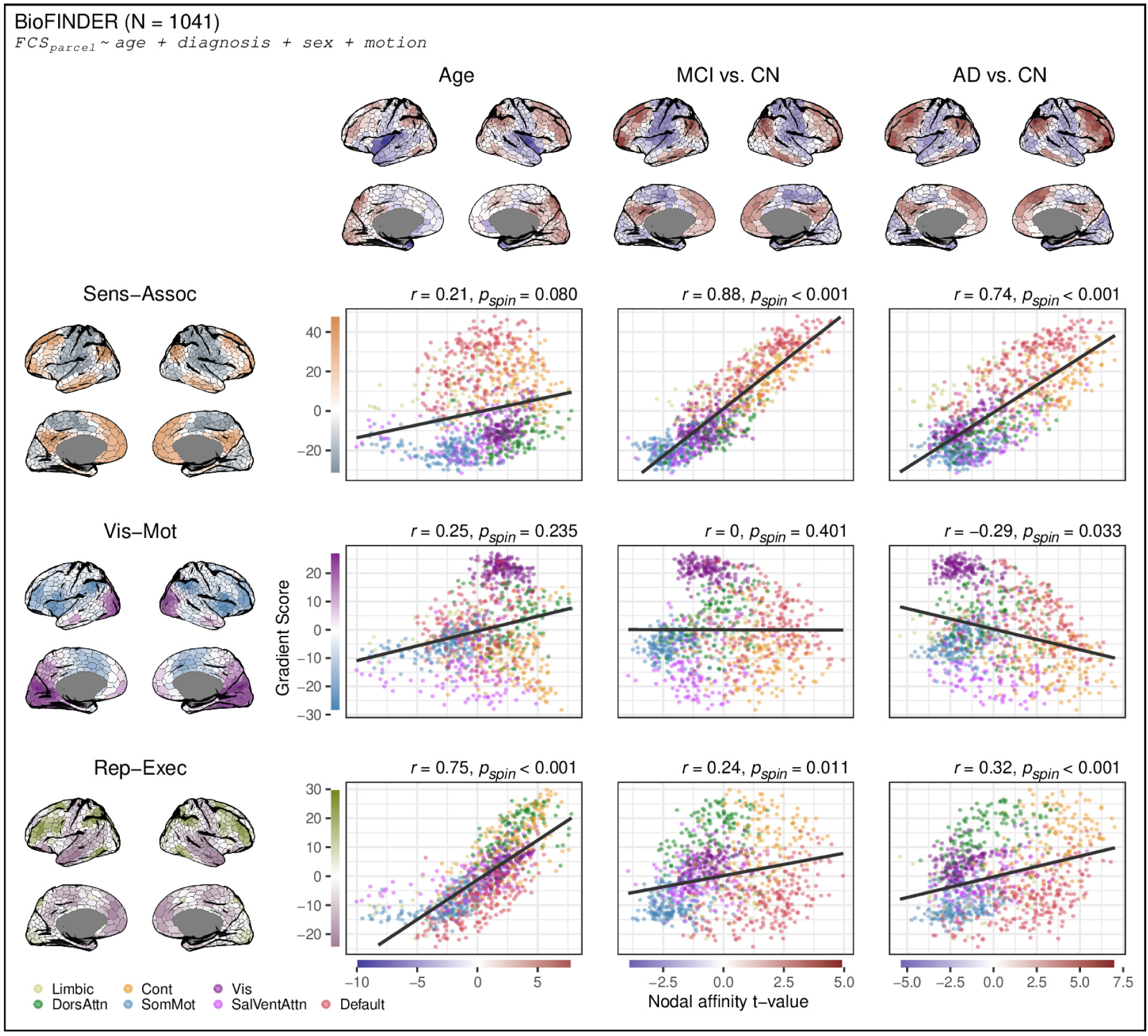
Clinical diagnosis reveals strong alignment with the sensory–association axis. Mild cognitive impairment and Alzheimer’s disease (vs cognitively normal) is associated with functional connectivity changes that closely follow the SA axis, paralleling effects observed for AD pathology, see Figure 2. Cortical maps display t-values from nodal linear models, and scatter plots show relationships between t-values and gradient scores, colored by network membership. Associations were quantified using Pearson correlation, with significance assessed using a spin test (one-sided, uncorrected; see Methods). The larger sample reflects greater availability of clinical diagnosis compared to pathology measures.

### Cross-sectional effects of age and AD pathology on FCS align with distinct gradients

In the BioFINDER-2 cohort, the spatial distribution of node-wise associations between FCS and AD pathology aligned with the SA axis (r = 0.74, p_spin_<0.001), reflecting decreases of FCS in sensory-motor (unimodal) regions and increases in associative (transmodal) regions with increasing pathology. Age effects were instead independently aligned with the RE axis (r = 0.75, p_spin_<0.001), with executive areas showing increasing and representational areas decreasing FCS with increasing age (Figure 2 A, Figure E3). Notably, age effects were not strongly aligned with the SA axis, and AD effects were not strongly aligned with RE axis, suggesting a double dissociation. All variance inflation factors were below 1.2, indicating no issue of multicollinearity between age and pathology. These findings were replicated using a subset of 129 participants from the ADNI with complete data (SA axis for pathology: r = 0.54, p_spin_<0.001; RE axis for age: r = 0.68, p_spin_<0.001; see Figure 2 B). We also ran the analysis with tau and Aβ separated, as well as with *APOE* ε4 status as a predictor. The results suggest that alignment to the SA axis is more closely related to tau than to Aβ positivity. In cognitively unimpaired individuals, Aβ positivity and APOE ε4 carriership showed modest inverse SA-alignment (Figure E4). Finally, in an additional model including cortical thickness, white-matter hyperintensity burden, lacunes, microbleeds, and CSF α-synuclein positivity (Figure E5), the characteristic gradient alignments for age (RE axis) and AD pathology (SA axis) remained largely unchanged, indicating that these effects are not explained by atrophy or cerebrovascular confounds. Interestingly, several of the additional covariates also showed independent effects aligning with the gradients.

**Figure 2:**
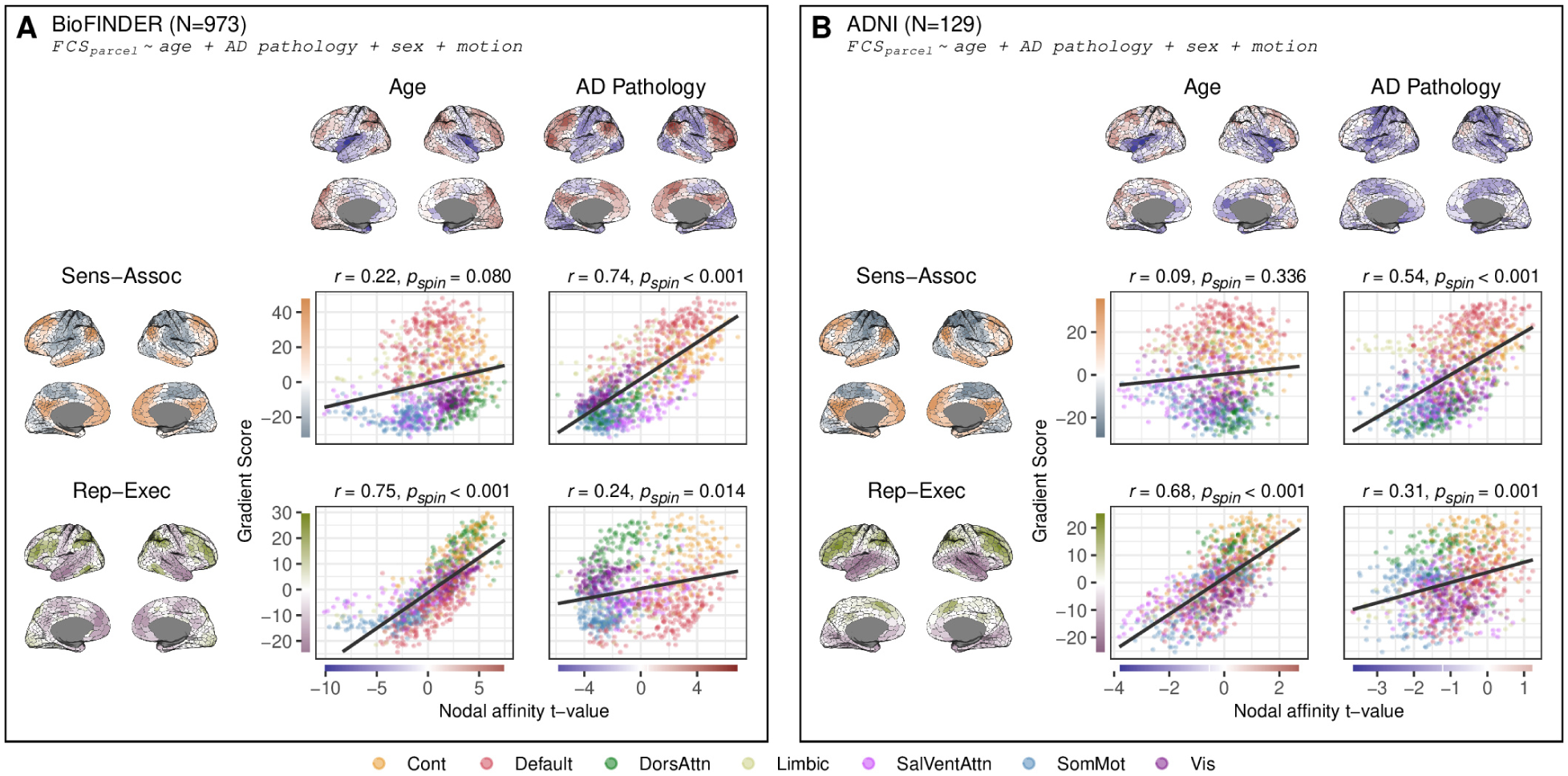
Age and AD pathology are associated with functional connectivity changes aligned with distinct organizational gradients. These dissociable alignments were observed in BioFINDER-2 (A), and in an external cohort, ADNI (B). Cortical maps display t-values from nodal linear regression models with nodal affinity as the outcome, using age and Alzheimer’s pathology as independent variables, covaried for sex and motion. Scatter plots show the relationship between t-values and gradient scores across nodes, colored by network membership. The relationship was quantified using Pearson correlation and significance assessed using a spin test (one-sided, uncorrected; see Methods). Color scales on the axes match the corresponding cortical t-maps. Points are colored by their Yeo network membership^38^. See Figure E3 for network delineation of the cortical maps of A.

**Figure E3:**
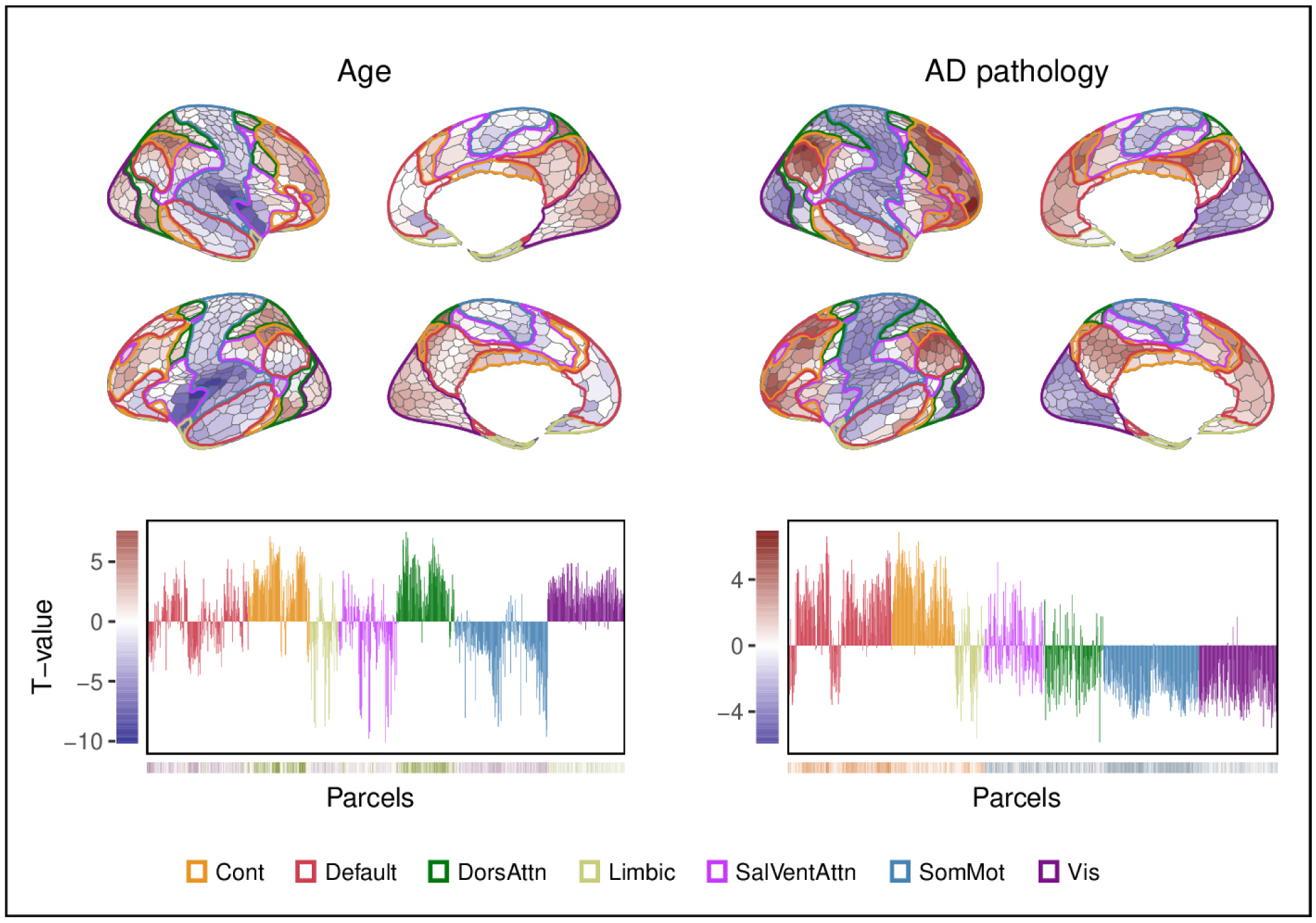
Yeo network distribution of gradient-aligned effects. Main results of Figure 2 A with the seven Yeo networks overlaid (top) and bar plots of the t-values from all parcels colored by their network membership (bottom). The colorbar on the y-axes of the histograms correspond to the color scale of the cortical maps, and the colorbar on the x-axes correspond to the color scale of the RE axis and the SA axis for Age and AD pathology, respectively.

**Figure E4:**
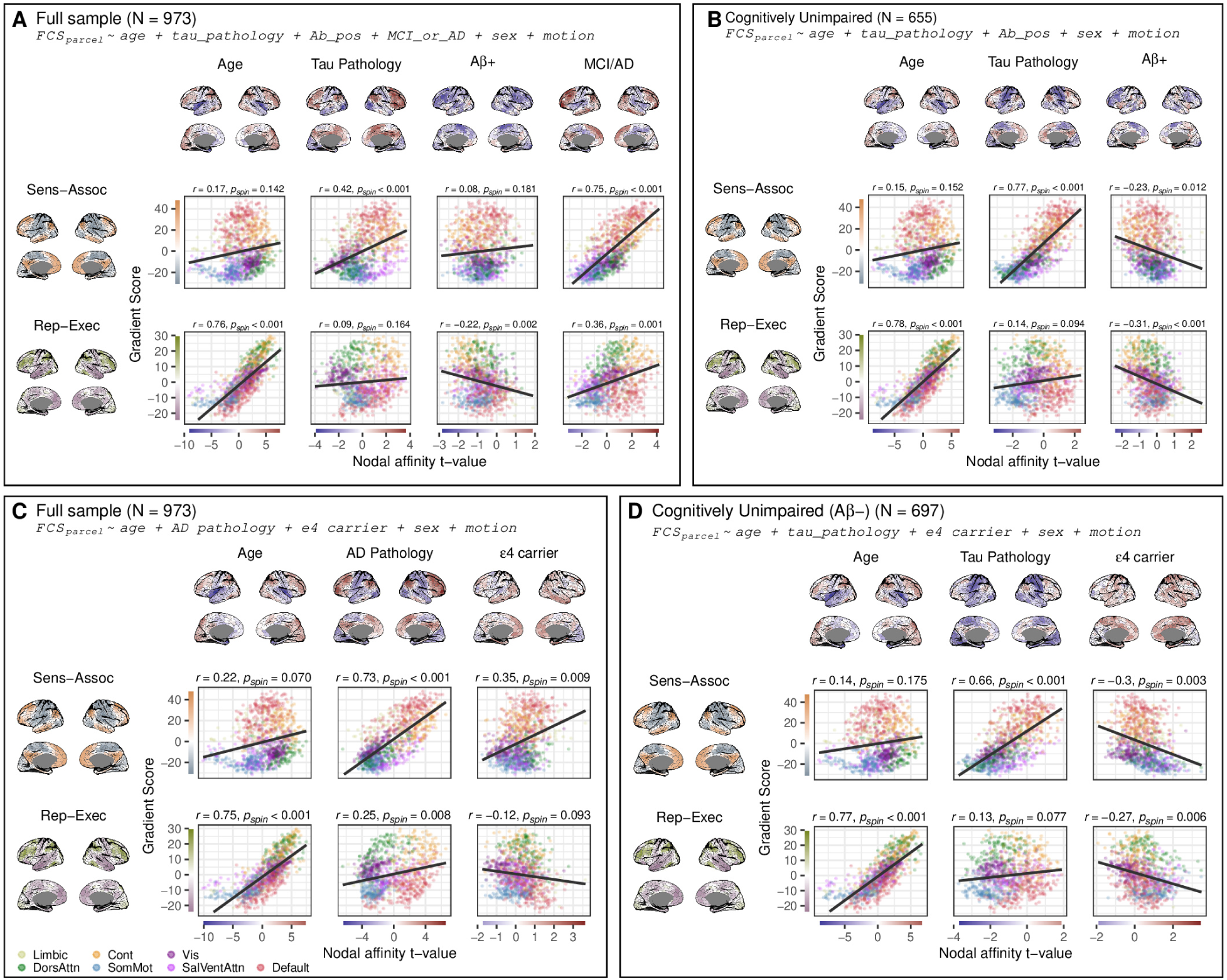
Gradient alignment is more related to tau than Aβ and APOE ε4 carriership has limited effects. A) Main results with separated pathology in the full sample controlling for diagnosis, due to the fact that all MCI/AD are Aβ+ by definition in this study. Tau pathology is calculated here as a non-linear mapping of tau PET SUVR from Braak12, Braak34, Braak56 in the same manner as for the main pathology composite. Aβ positivity was determined using previously established amyloid-PET cut-offs derived in^39^. B) Main results with separated pathology in cognitively unimpaired. Tau effects on FCS show considerably stronger alignment with the sensory-association axis compared to Aβ positivity. C) Main result with *APOE* ε4 carriership in the model in the full sample. D) Since the relationship between being an ε4 carrier and being Aβ+ is strong, and being Aβ+ is strongly related to being MCI/AD, we also ran the analysis in cognitively unimpaired Aβ- individuals. Here, we can see that being an ε4 carrier in itself does not give rise to any marked gradient alignment.

**Figure E5:**
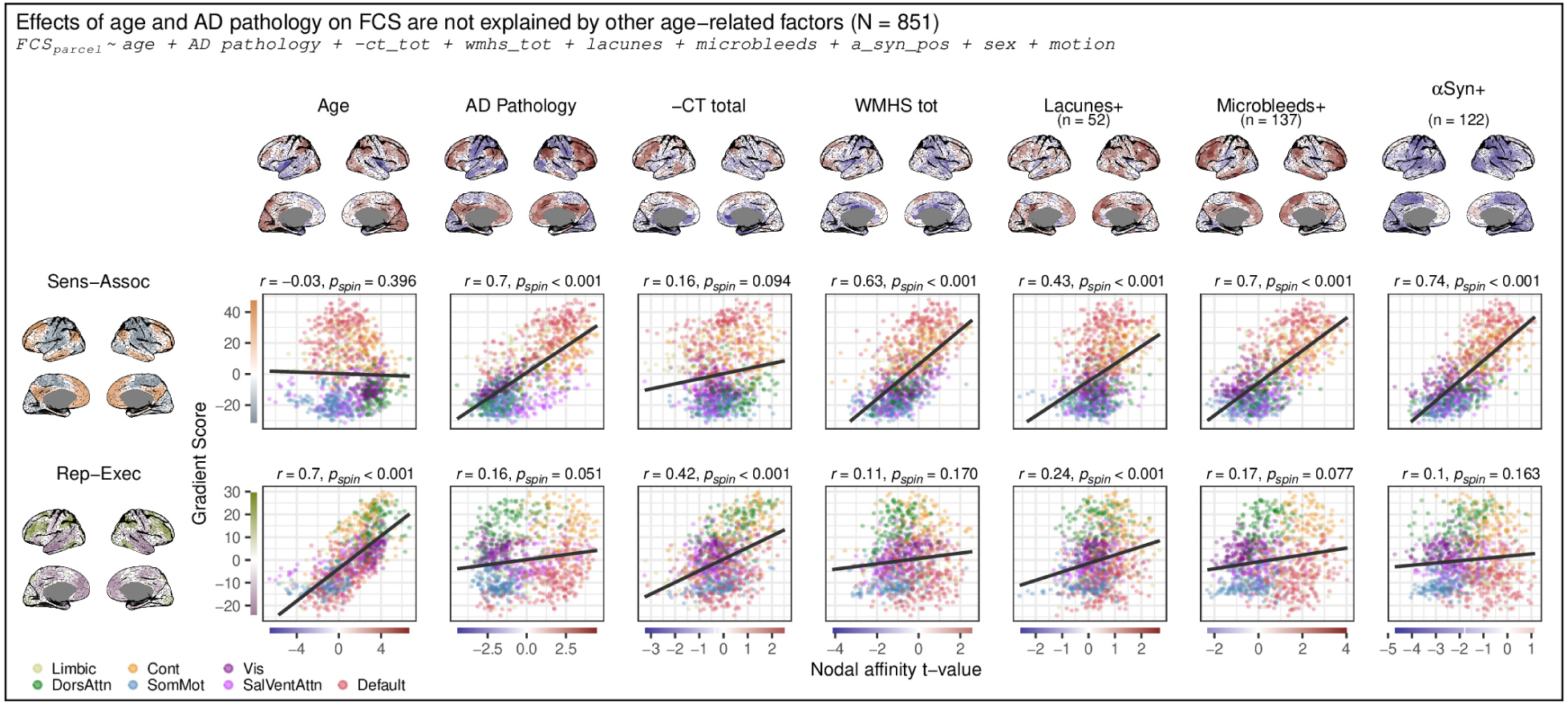
The FCS patterns associated with age and AD remained stable after including atrophy and several vascular markers as covariates in the main analysis. The parcel-wise model included age, tau pathology, cortical thickness in the AD atrophy signature region, white-matter hyperintensity burden, presence of lacunes, presence of microbleeds, and CSF α-synuclein positivity (RTQuIC). Scatterplots show the spatial correlation between each effect map and the sensory–association and representational–executive axes. The principal age- and tau-related gradient alignments reported in the main manuscript remain robust after accounting for these factors, which themselves seemingly give rise to gradient-aligned effects.

### FCS effects and their alignment with gradients vary along the age- and pathology spectra

Prior studies on FC in AD have reported an initial pattern of regional hyperconnectivity, which is subsequently followed by connectivity declines^1,19,40^. Nonlinear relationships between age and FC have also been reported^8,41^. We therefore sought to investigate whether these nonlinearities were reflected similarly in gradient-like FCS changes. To investigate these potential nonlinearities, generalized additive models (GAMs) were fitted with FCS at each parcel as the outcome, and age, pathology and relevant covariates as predictors. The derivatives of these models capture the direction and strength (i.e., slope) of each predictor’s effect across its range. For each predictor (age and pathology), the estimated slopes result in cortical “slope-maps” that reflect the regional strength and direction of the predictor’s effect, at any value of the predictor. This allows us to observe increases or decreases at specific points across the age and pathology spectra. For pathology, the relationship between the SA axis and the slope-maps increased up to a pathology level of ∼0.25 (representing early AD pathology), remaining relatively high until ∼0.5 (representing middle-stages of AD pathology), after which it declined sharply with further accumulation (Figure 3 B.2). For age, the relationship between the RE axis and the slope-maps remained high between ages 55 and 70, but essentially vanished after age 80. These results suggest that gradient-like FCS patterns vary systematically across different levels of age and pathology, and appear most pronounced at intermediate levels of each factor.

**Figure 3:**
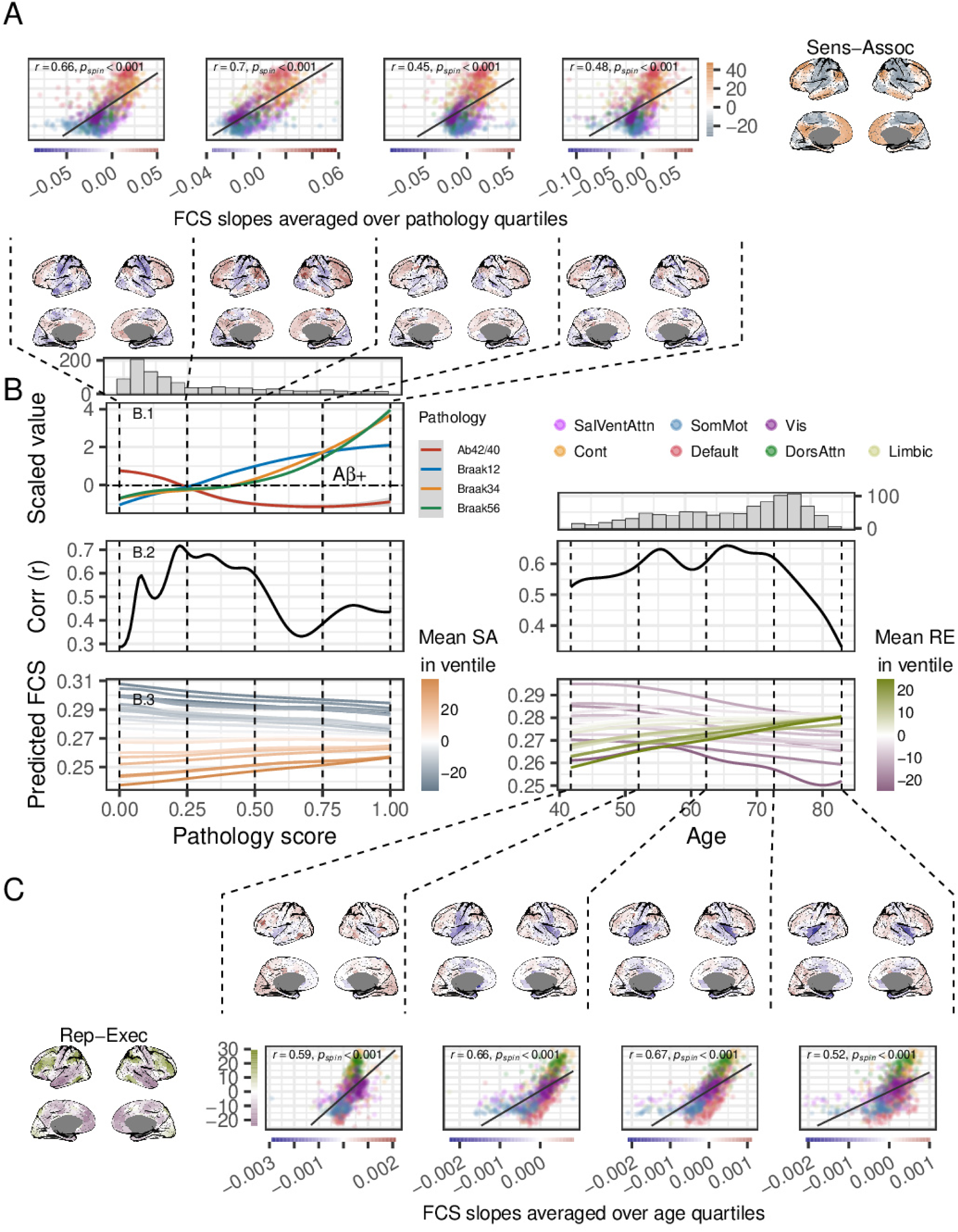
Nonlinear models reveal that the alignment of age- and pathology-related FCS effects with organizational gradients varies across the age and AD pathology continuum. Generalized additive models (GAMs) were fitted for each parcel, with nodal affinity as the outcome, and age and AD pathology as (non-linear) smooth terms, while controlling for sex and motion. Predicted trajectories and their derivatives were calculated. (A): Top row: Scatter plots of averaged slope coefficients across pathology quartiles vs. SA axis scores, with parcels colored by network membership. Bottom row: Cortical maps of averaged slope coefficients. (B.1): Nonlinear relationship (LOESS curves with 95% CI) between the composite AD pathology score and scaled pathology measures. (B.2): Correlation coefficients showing how alignment between trajectory derivatives and gradient scores changes across pathology (SA axis) and age (RE axis). (B.3): Predicted nodal affinity trajectories for pathology and age, averaged within gradient-based ventiles (20 equally sized groups), colored by mean gradient scores. Marginal histograms in B shows the sample sizes across pathology score and age from the original estimation. (C): Same as A, but for age quartiles.

Regional FCS trajectories, averaged within 20 quantile bins (ventiles) based on gradient values, are shown in Figure 3 B.3. Across the pathology spectrum, average FC in transmodal ventiles increased relatively linearly, while FCS in unimodal ventiles decreased at a similar rate. Sensory-motor (unimodal) ventiles consistently exhibited higher FC than associative (transmodal). Across age, FCS increased in executive ventiles and decreased in representation ventiles. While executive ventiles had lower FCS than representation regions in younger age, this pattern reversed in older age.

Prior work has suggested that both phenotypic heterogeneity and age–pathology interactions may influence large-scale functional organization in AD^31^, raising the possibility that these factors may contribute to the decrease in alignment we see over progressive accumulation of pathology. To examine this, we first included an age × pathology interaction term in the parcel-wise linear models. This interaction showed an inverse relationship with the sensory–association axis, indicating that increasing age attenuates the gradient-aligned FC effects associated with pathology. Consistent with this, when stratifying patients by age (<65 vs ≥65 years) and repeating the nonlinear analyses from Figure 3, the older group showed a steeper decline in gradient alignment with increasing pathology. However, a reduction in alignment was also observed in the younger group, indicating that age–pathology interactions alone do not explain the loss of alignment at higher pathology levels. In contrast, stratifying patients into amnestic and non-amnestic groups based on cognitive domain scores revealed similar alignment trajectories, with no clear differences between groups (see Figure S2).

### Longitudinal increase in pathology is associated with gradient-aligned FCS changes

A subset of participants in the BioFINDER-2 sample (N = 378) had longitudinal data available for all variables except CSF Aβ (see Table 2 for longitudinal characteristics, number of visits etc.). To determine whether the gradient-aligned FCS changes from previous analyses reflect within-subject responses to pathology (as opposed to cross-sectional associations), we fit linear mixed-effects models. Since longitudinal CSF Aβ data were unavailable, the pathology score was re-calculated using only tau PET from Braak I-II, III-IV and V-VI regions of interest. The baseline effects in this sample (representing between subject differences; Figure 4 A) closely mirrored the previous cross-sectional findings (Figure 4 A). The longitudinal change in pathology (ΔPathology, representing within-subject changes) also showed gradient-aligned FCS changes (SA: r = 0.59, p_spin_<0.001), confirming that within-subject pathology accumulation is associated with within-subject FCS increases and decreases along organisational axes. Interestingly, there was also a modest relationship between the RE axis (r = 0.38, p_spin_<0.001) and longitudinal change in pathology.

**Figure 4:**
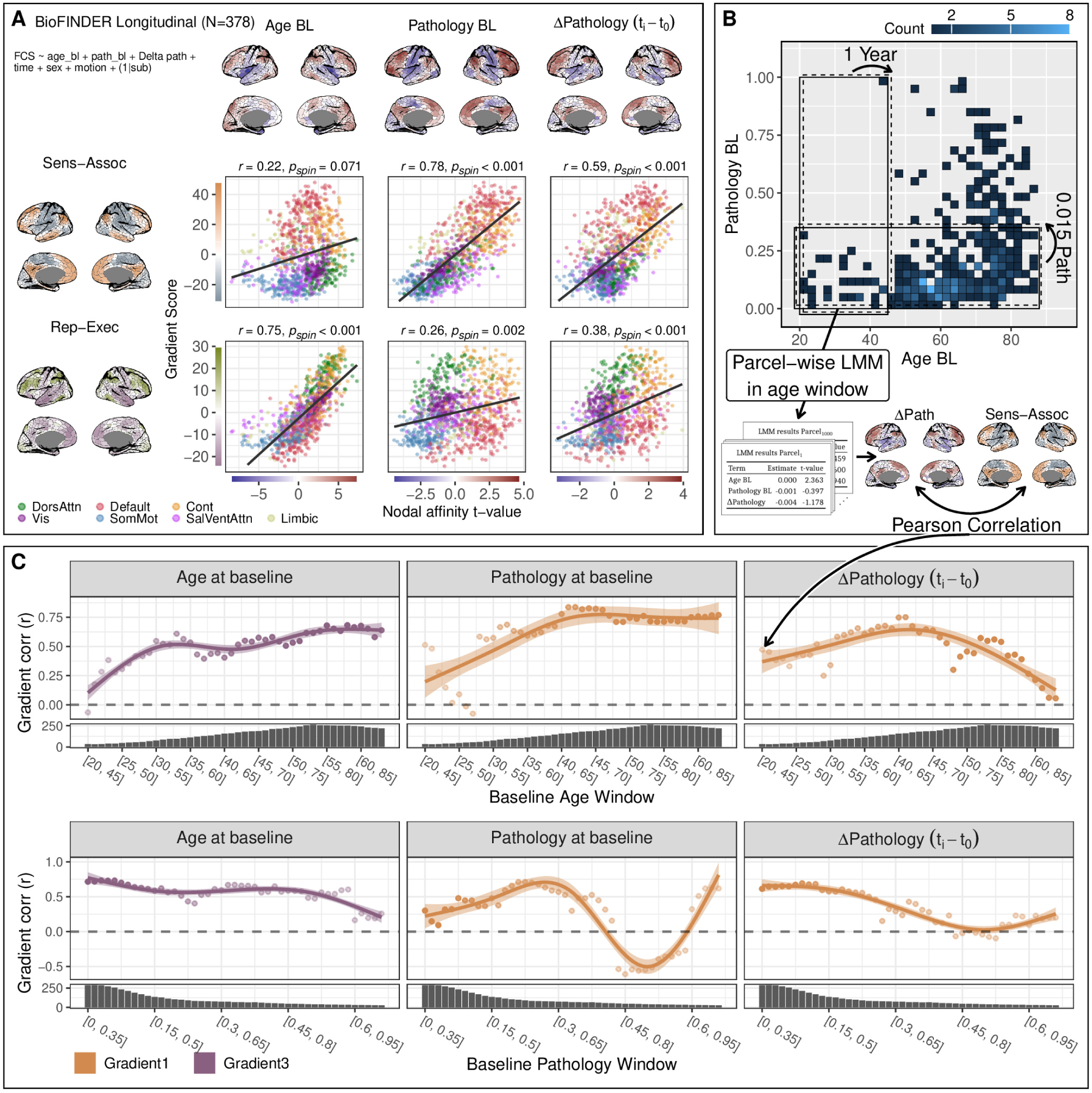
Within-subject change in pathology is related to gradient-aligned FCS effects. A: A linear mixed-effects model was applied at each parcel, with FC as the outcome and baseline age, baseline pathology (tau PET SUVR from Braak I-VI), Δpathology (longitudinal difference in tau PET from baseline) as fixed effects and random intercepts for subjects. Cortical maps display t-values from the parcel-wise models, while scatter plots show the relationship between t-values and gradient scores, colored by network membership. The relationship was quantified using Pearson correlation and significance assessed using spin tests (one-sided, uncorrected; see Methods). B: Schematic of the sliding window approach used to assess longitudinal nonlinearities in the relationship between FCS and gradients across the aging and pathology spectra. The heatmap represents a bivariate histogram. Windows are represented as “moving” rectangles. Age windows spanned 25 years and were incremented by 1 year, while pathology windows spanned 0.35 pathology units and were incremented by 0.015 (indicated by the curved arrow), resulting in 44 age windows and 45 pathology windows, all having ≥ 25 subjects. For each window we fit parcel-wise linear mixed models, t-maps for each fixed effect of interest were extracted and the t-maps correlated (Pearson correlation) with the SA and RE axes. C: Results of the sliding window analysis showing the dynamic relationship of AD pathology on FC gradient alignment. The correlation values on the scatters are overlaid with an estimation from generalized additive models, with 95% confidence intervals. For the age terms in each windowing, correlations are shown between the t-map and the RE axis, while for the pathology terms, they are shown be-tween the t-maps and the SA axis. Marginal plots show the sample size for each window. Results from the age windowing show that baseline age and pathology effects on the RE and SA axes, respectively, are mostly stable across the adult lifespan. Across the pathology windows, baseline pathology effects increase in gradient alignment over the initial windows after which they decrease. This pattern was mirrored in Δ pathology, but the within subject change preceded the baseline effects with peaks and declines in earlier windows.

**Table 2:**
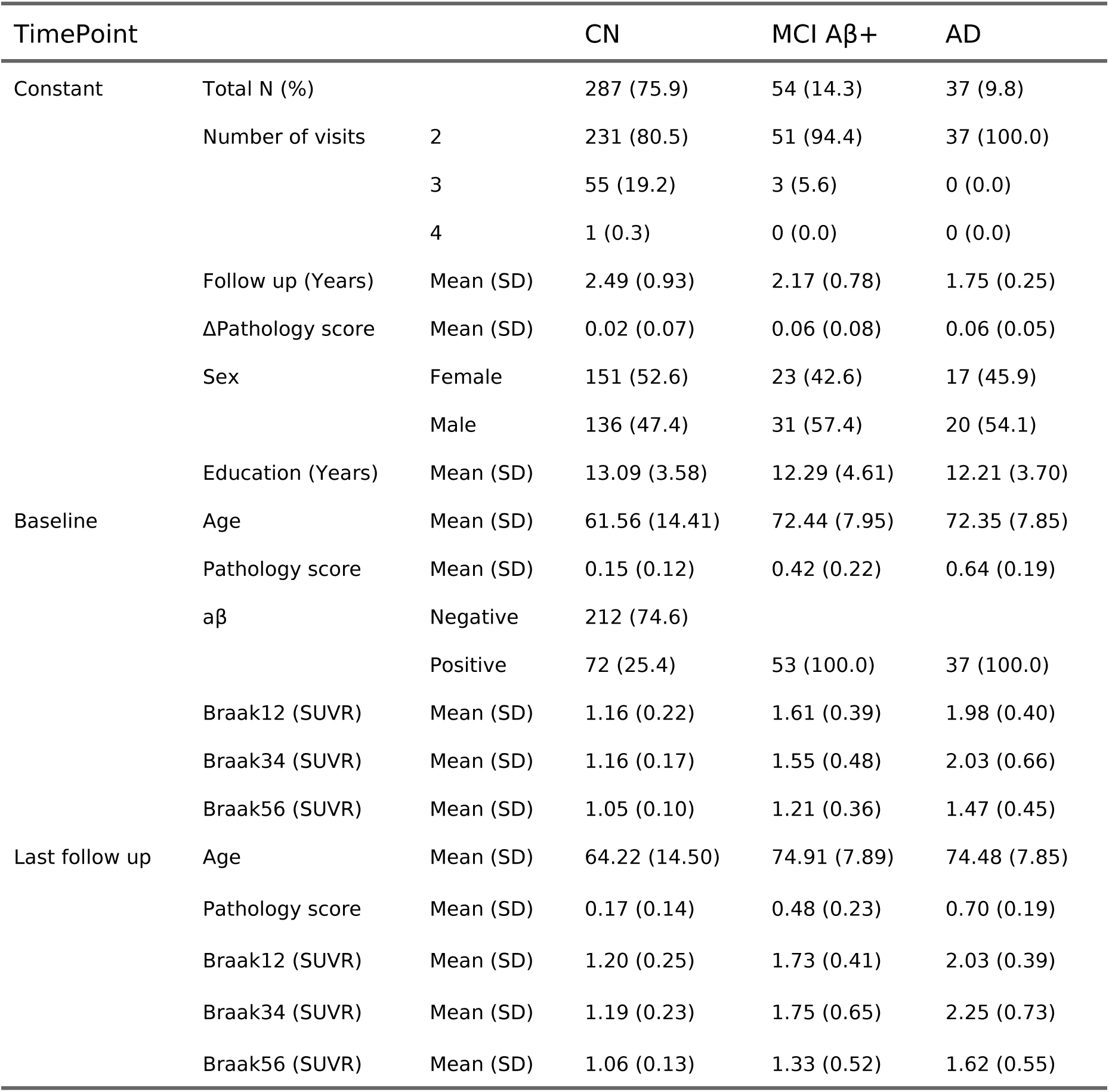
Characteristics of the longitudinal BioFINDER-2 sample. Because no CSF Aβ data were available longitudinally for AD dementia patients, pathology scores were recomputed using only tau PET SUVR from Braak I-II, III-IV, V-VI regions. TimePoint indicates whether a variable is baseline, last available follow-up, or consistent across time. Abbreviations: FD = frame displacement (scanner motion); SD = standard deviation, CN = cognitively normal; MCI = mild cognitive impairment; AD = Alzheimer’s disease.

**Table 3:**
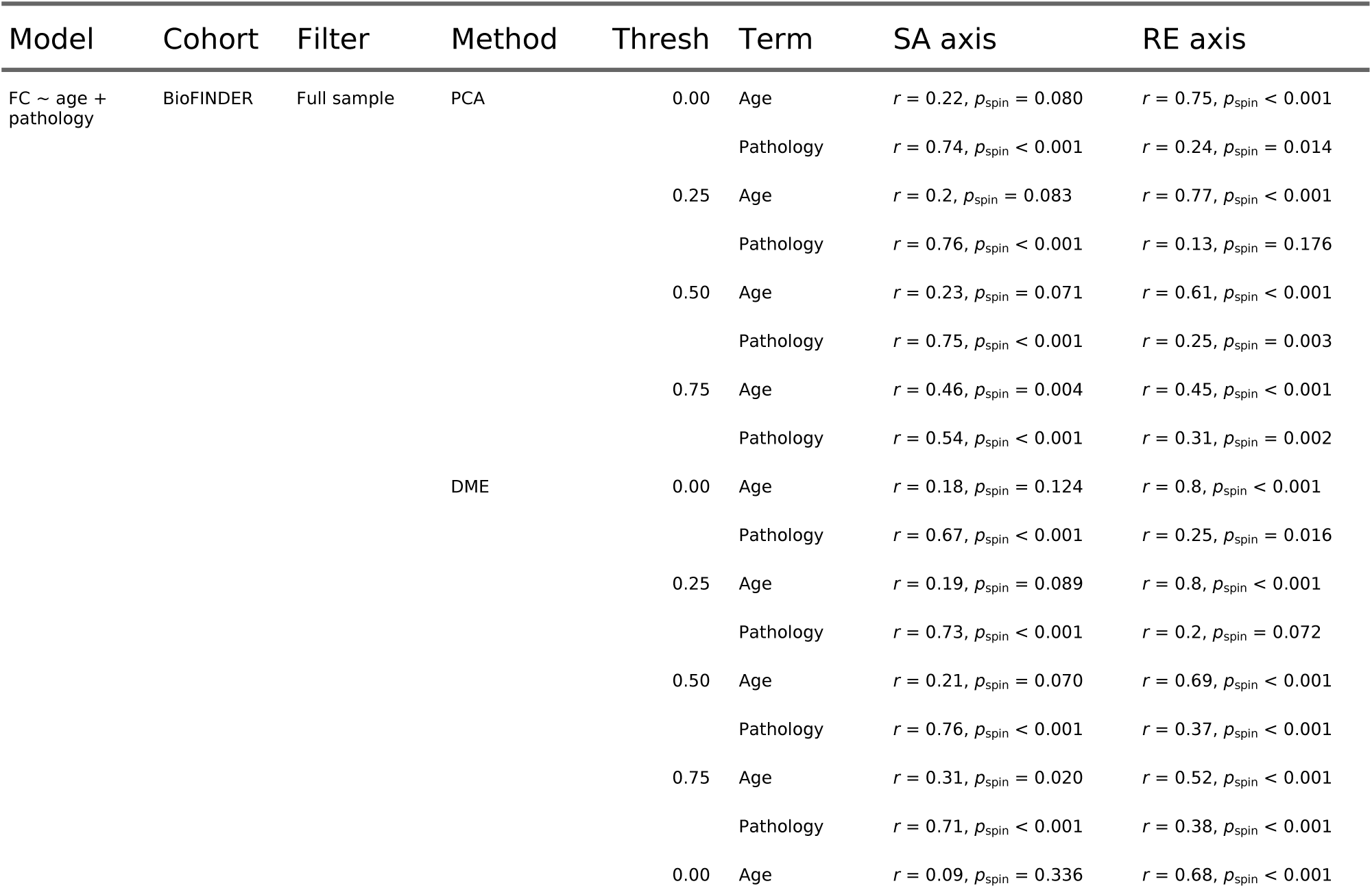
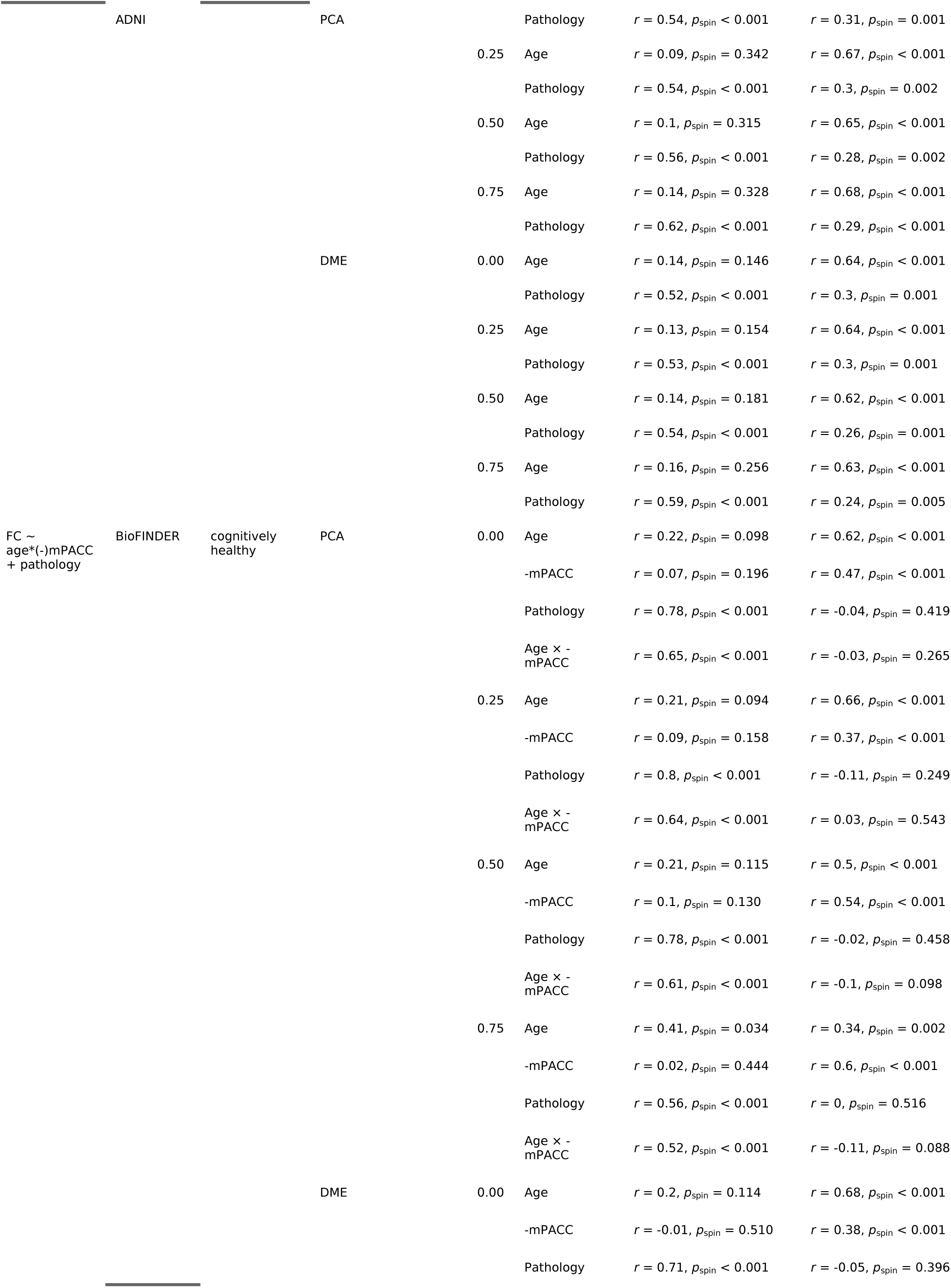
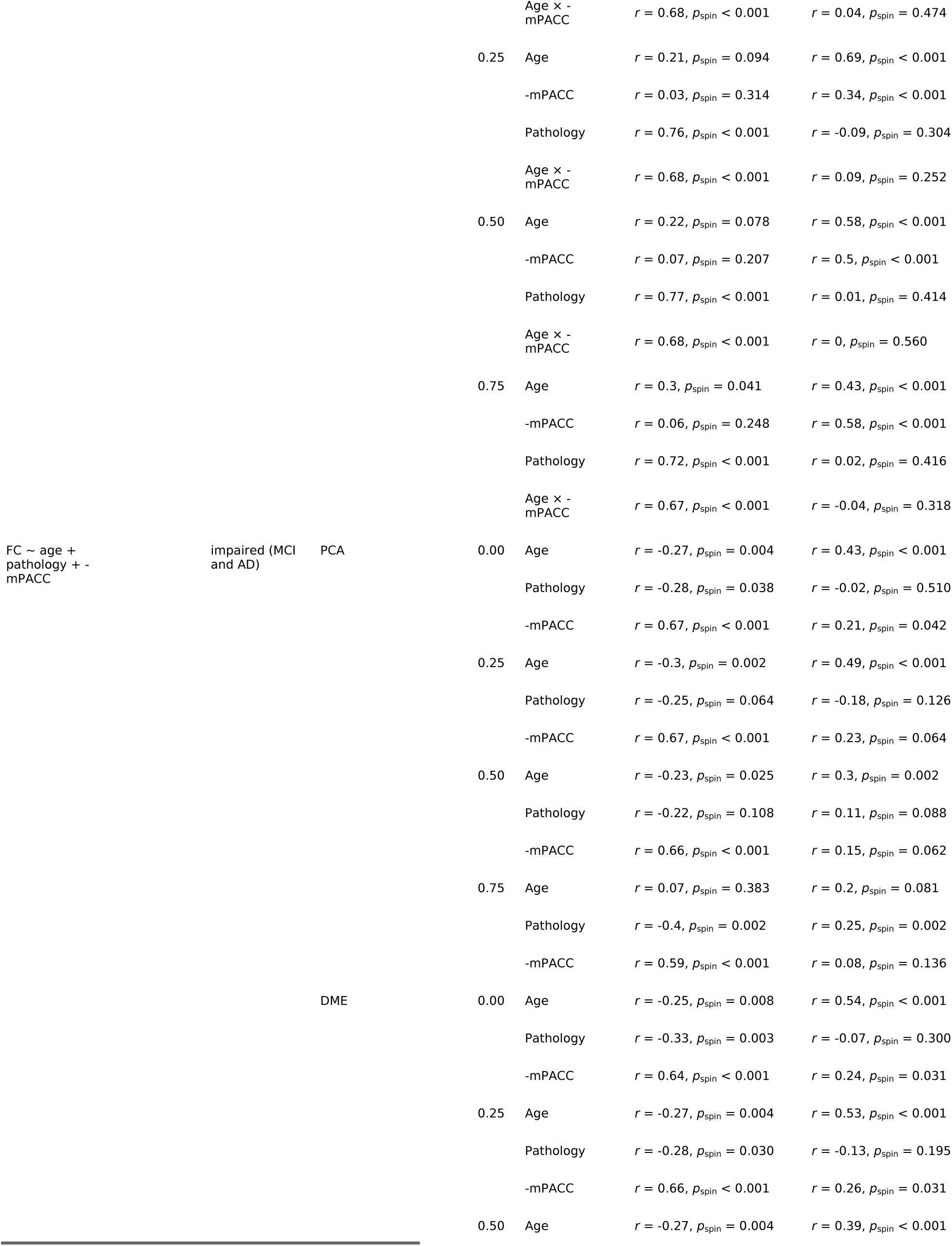
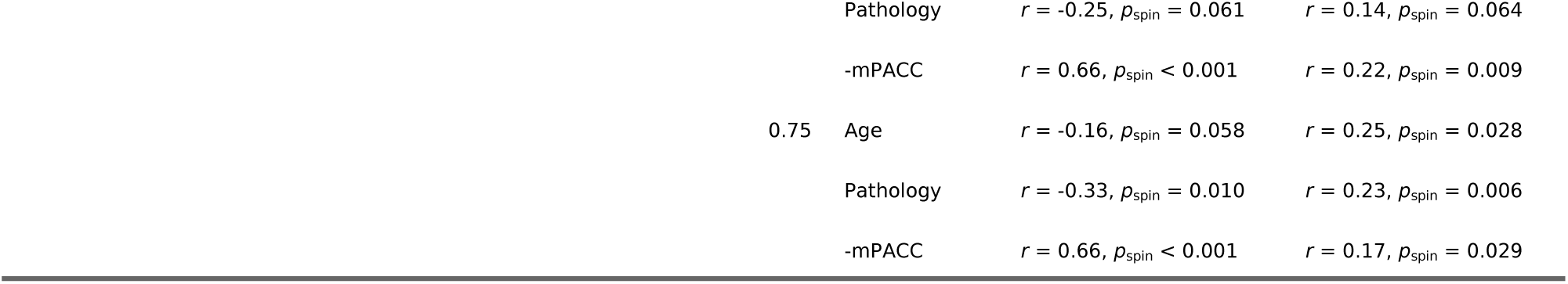
Gradient derivation method and FC thresholding has limited effect on the main results. This table shows the main analyses from Figure 2 and Figure 5 using gradients derived with principal component analysis (PCA) or diffusion map embedding (DME) over four different thresholding values: 0.00, 0.25, 0.50 and 0.75. A threshold of 0.75 means that 75% of the lowest values in the connectivity matrix is zeroed out. In the columns SA axis and RE axis, the Pearson correlation (r) is shown with accompanying p-value from spin tests (one-sided; see Methods), quantifying the relationship between the t-values for each specific term and the gradient scores based on the derivation method and thresholding specified in those two columns. Gradients were derived in all analyses from the average connectome of individuals without cognitive impairment, apoe4 non-carriers without abnormal amyloid levels. In BioFINDER only individuals ≤60 years of age were included while in ADNI the whole age range was used due to the smaller sample size. For the DME method this matrix is then converted into an affinity matrix by taking the cosine similarity between each pair of connectivity profiles, the components are then calculated as outlined in^20^.

To explore whether the previously identified nonlinearities were also reflected longitudinally, we employed a sliding window approach, examining how gradient alignment varies across different baseline ages and baseline pathology levels. Figure 4 B provides a methodological overview, and Figure 4 C displays the gradient alignment (correlation values) for each window and term. For clarity, we only present the relevant gradient correlations for each term as indicated by our prior cross-sectional analyses, though Figure S3 provides both. Age-windowing revealed relatively strong RE correlations for baseline age across age windows above ∼30 years of age, suggesting that a 25-year interval is long enough to capture this effect across most of the adult lifespan). Baseline pathology effects remained consistently aligned with the SA axis across all age windows beyond 40 years of age. The within-subject pathology change (ΔPathology) showed the strongest SA alignment at approximately the same age window as baseline pathology, but decreased somewhat in older age.

Across the pathology windows, baseline age showed a decreasing association with the RE axis, with weaker correlations in windows containing higher baseline pathology. Baseline pathology effects, in contrast, initially increased in alignment with the SA axis before declining and even showing negative correlations (indicating reversal from prior states) at higher pathology levels. This peak-and-decline pattern echoes the findings from the cross-sectional GAM analysis. Interestingly, ΔPathology followed a similar trajectory but preceded the baseline pathology effect, peaking and declining in earlier pathology windows. While we observed an increase at the highest end of the baseline pathology windows, we also note limited sample size and reduced reliability of estimates at distributional extremes. Overall, these findings corroborate and extend our cross-sectional results, demonstrating that AD pathology-related FCS changes have a temporal dynamic. Within-subject increases in tau pathology are initially associated with FCS changes along the SA axis, but this relationship weakens as baseline pathology advances. Moreover, the stronger ΔPathology effects at lower pathology windows, compared to baseline pathology, suggest that gradient-aligned connectivity changes may be detectable at very early stages of pathology.

### Gradient-aligned FCS patterns reflect cognitive status

In the previous analyses, we linked gradient-like connectivity alterations to two factors known to affect cognition: age and AD. However, these analyses did not directly examine the relationship between connectivity and cognitive performance, which is essential for understanding the implications of these findings. Hence, to assess the relationship between cognition and FCS alterations in our presented framework we examined the association between FCS and cognitive performance in two groups: cognitively healthy, CSF Aβ42/40-negative, *APOE* ε4 non-carriers (N = 310) and individuals with mild cognitive impairment (MCI) or AD dementia (N = 258). This split was intended to assess whether the FCS patterns are related to cognition in the absence of cognitive decline attributable to AD dementia. Cognitive performance was measured using a modified Preclinical Alzheimer Cognitive Composite (mPACC) score^42^, zero-normalized to a healthy control group over 60 (see Methods for details).

In the healthy group, node-wise linear regression was conducted with mean centered age, (inverted) mPACC and their interaction as predictors. Due to common age-related tau accumulation^43^, we adjusted for pathology load in the healthy group (see Figure S4 for analyses without this adjustment and distribution of Tau SUVR within this group). Results (Figure 5 A) confirmed alignment of age effects with the RE axis (r = 0.62, p_spin_<0.001). RE-like changes in FC were also associated with worse cognition (r = 0.47, p_spin_<0.001). This means that, at the sample mean age (62 years), people with poorer cognition generally have decreases of FCS in executive regions and less decreases (or increasing) FCS in representation regions (i.e. have more of a RE-like FCS pattern). This main effect was also present in the absence of the interaction term (Figure S5). The interaction between age and cognition, however, showed an SA-like pattern (r = 0.65, p_spin_<0.001) – meaning that older people with poor cognition generally have decreased FCS in sensory-motor regions and less decreased (or increased) FCS in associative regions (i.e., have more of an SA-like FCS pattern). Surprisingly, we found that the AD pathology (in this case, early stage tau pathology, since all subjects in this group are Aβ-) in the cognitively healthy group also aligned with the SA axis (r = 0.78, p_spin_<0.001), again suggesting that gradient-aligned FCS changes may indicate emerging pathology.

**Figure 5:**
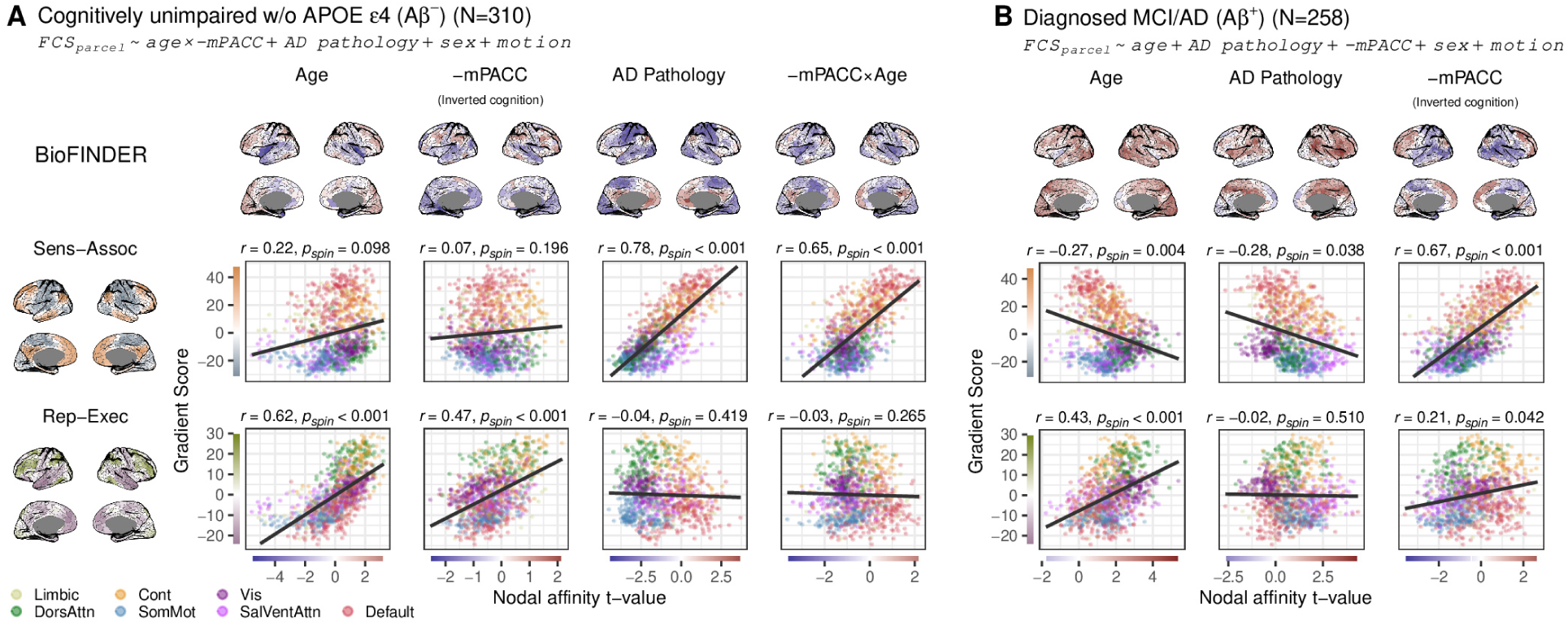
Cognitive status modifies the relationship between AD pathology, age, and FCS changes along the SA axis. Cross-sectional results are shown for (A) cognitively unimpaired Aβ-, *APOE* ε4 non-carriers and (B) cognitively im-paired individuals with mild cognitive impairment (MCI) or AD dementia. In A, models include age, pathology, cognition (mPACC), and an cognition × age interaction (with centered predictors and inverted cognition scores for interpretability). In B, models include age, pathology, and cognition without interactions. Tau-driven AD pathology in the cognitively unimpaired group is associated with SA-like FCS changes whereas it is absent in the MCI/AD dementia group. Instead, worse cognition (-mPACC in B) in the MCI/AD dementia group is associated with SA-like changes, which closely resembles the effects seen in older cognitively unimpaired individuals (-mPACC x Age in A).

In the cognitively impaired group, parcel-wise regression was performed to examine FCS associations with AD pathology and (inverted) mPACC, adjusted for age. Surprisingly, the spatial distribution of AD pathology effects no longer resembled an SA-like pattern in this group (see Figure S6 for this analysis with gradients derived from the full sample). Instead, an SA-like pattern emerged for worse cognition, regardless of pathology load or age (Figure 5 B, see also Figure S7 for the same subgroup analysis but without cognition). This suggests that in the clinical phase of AD, cognitive difficulties co-occur more strongly with SA-aligned FCS changes than the connectivity changes related to underlying pathology. Supplementary analyses (Figure S8) confirmed that adding an interaction between pathology and cognition did not alter this main effect.

### Gradient-aligned FCS and domain-specific cognitive performance

Prior work has shown that aging and Alzheimer’s disease disproportionately affect different cognitive domains, with aging more strongly impacting executive function and AD pathology more strongly impacting memory^44,45^. This raises the possibility that the dissociable gradient alignments observed for age and pathology reflect differences in the cognitive pressures exerted by these processes. To examine this, we derived domain-specific cognitive scores (global, memory, executive, language, visuospatial) using a bifactor model estimated on cognitively healthy participants below 70 years (n=285, see Figure 6 A and Methods). Consistent with prior literature, aging was more strongly associated with a decrease in executive performance, whereas AD pathology was more strongly associated with decreased memory performance in this cohort (see Figure 6 C). We then included these domain scores, controlling for global cognition, in parcel-wise linear models. Global cognition showed a strong inverse alignment with the SA axis, whereas both memory and executive performance showed positive alignment with both the SA and RE axes (Figure 6), with good memory and executive performance having stronger alignment with the SA and RE axis respectively, indicating that higher domain-specific performance was generally associated with greater gradient alignment. To further assess whether differential cognitive pressures contribute to the age- and pathology-related FCS patterns, we conducted parcel-wise mediation analyses (Figure 6 D). In general, the indirect effects of AD pathology on memory performance through FCS tended to be positive, i.e. the pathology effects on FCS did not seem to be detrimental for memory performance. A similar, but less clear, pattern emerged for the indirect effects of age on executive performance. Interestingly, seemingly indirect effects beneficial for one domain tended to be detrimental for the other, both for aging and AD pathology (Figure 6 D, bottom row). It should of course be noted that these indirect effects are small and none would likely survive multiple comparisons correction.

**Figure 6:**
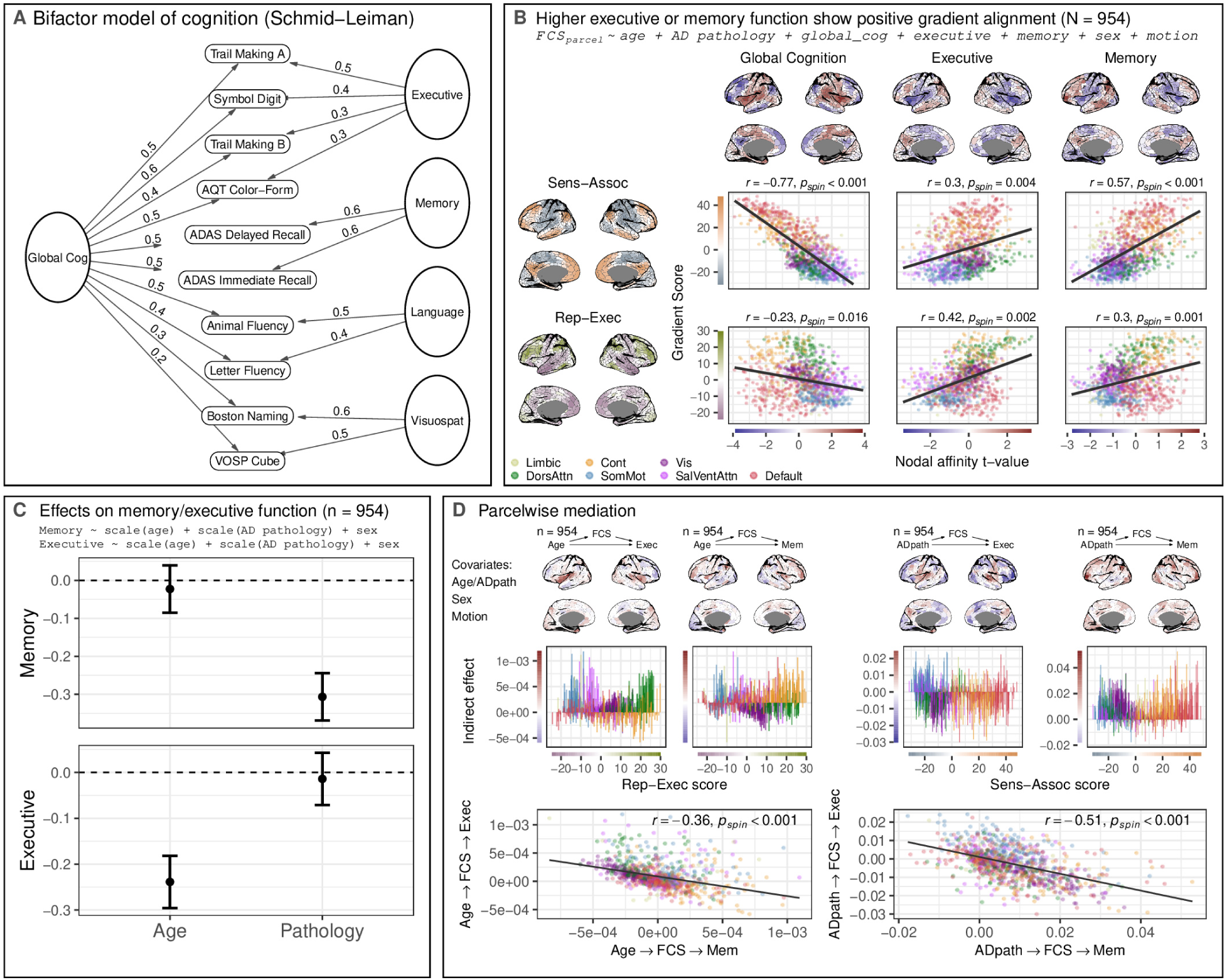
Domain-specific cognition maps onto functional connectivity gradients that differentially relate to aging and Alzheimer’s disease. (A) Bifactor (Schmid–Leiman) cognitive model. Ten cognitive tests were decomposed into a general factor (“g”) and four orthogonal domain factors (memory, executive, visuospatial, language; see Methods). Standardized loadings are shown for each test on both the general and domain-specific factors. (B) parcel-wise associations between domain-specific cognition and FCS. Parcel-wise linear models were fit with FCS as the outcome and domain scores as predictors, covarying for age, AD pathology, sex, motion, and global cognition. Higher scores on the domain-specific factors were associated with positive gradient alignment. Scatterplots show spatial correlations between parcel-wise t-values and the sensory–association and representational–executive axes. (C) Effects of age and AD pathology on cognitive domains. Regression coefficients (±95% CI) from models predicting domain-specific factor scores indicate that age is more strongly associated with executive decline, whereas AD pathology is more strongly associated with memory decline. The sample (N = 954) consists of all individuals from the original sample with values on at least 5 of the 10 cognitive tests. (D) Parcel-wise mediation analyses. Indirect effects quantify whether age- or pathology-related changes in FCS mediate their effects on executive and memory performance. Domain scores were orthogonalized with respect to global cognition, and models co-varied for pathology in age analyses and for age in pathology analyses. In the middle row indirect effects are visualised across their position on the two respective axes (RE for age and SA for pathology). Positive effects indicate beneficial mediation and negative effects detrimental mediation. The negative correlation (bottom row) between the indirect effects for the two different domains indicates that effects seemingly beneficial for one domain may be detrimental for the other.

### Sensitivity analyses

To demonstrate the robustness of our results to methodological choices, we performed several sensitivity analyses. First, we derived functional gradients using two different dimensionality reduction techniques for gradient-estimation and across four correlation-coefficient thresholds for each main analysis (32 replications in total), and found that the alignment with the t-maps across approaches were consistent (Table S1). We also correlated the t-maps within the BioFINDER sample to the gradients derived from ADNI and vice versa with comparable results (Figure S9 A,B). Interestingly, we repeated the main analysis using nodal connectivity strength (the average connection weight of each parcel and more conventional FC measure) instead of nodal affinity and observed comparable spatial patterns (Figure E6). We further replicated the results using within-network affinity (Figure S10), between-network affinity (Figure S11), and nodal affinity computed directly from unthresholded Pearson correlations (Figure S12), also with similar spatial patterns. Additionally, we ran the main analysis with a pathology score calculated after pooling BioFINDER and ADNI rather than computing the pathology scores separately, and the results did not deviate to any great extent from the original (Figure S9 C,D). Parcel-specific tau PET models produced results comparable to the global tau models, suggesting that regional tau levels mainly reflect global tau levels in this context (Figure E7).

**Figure E6:**
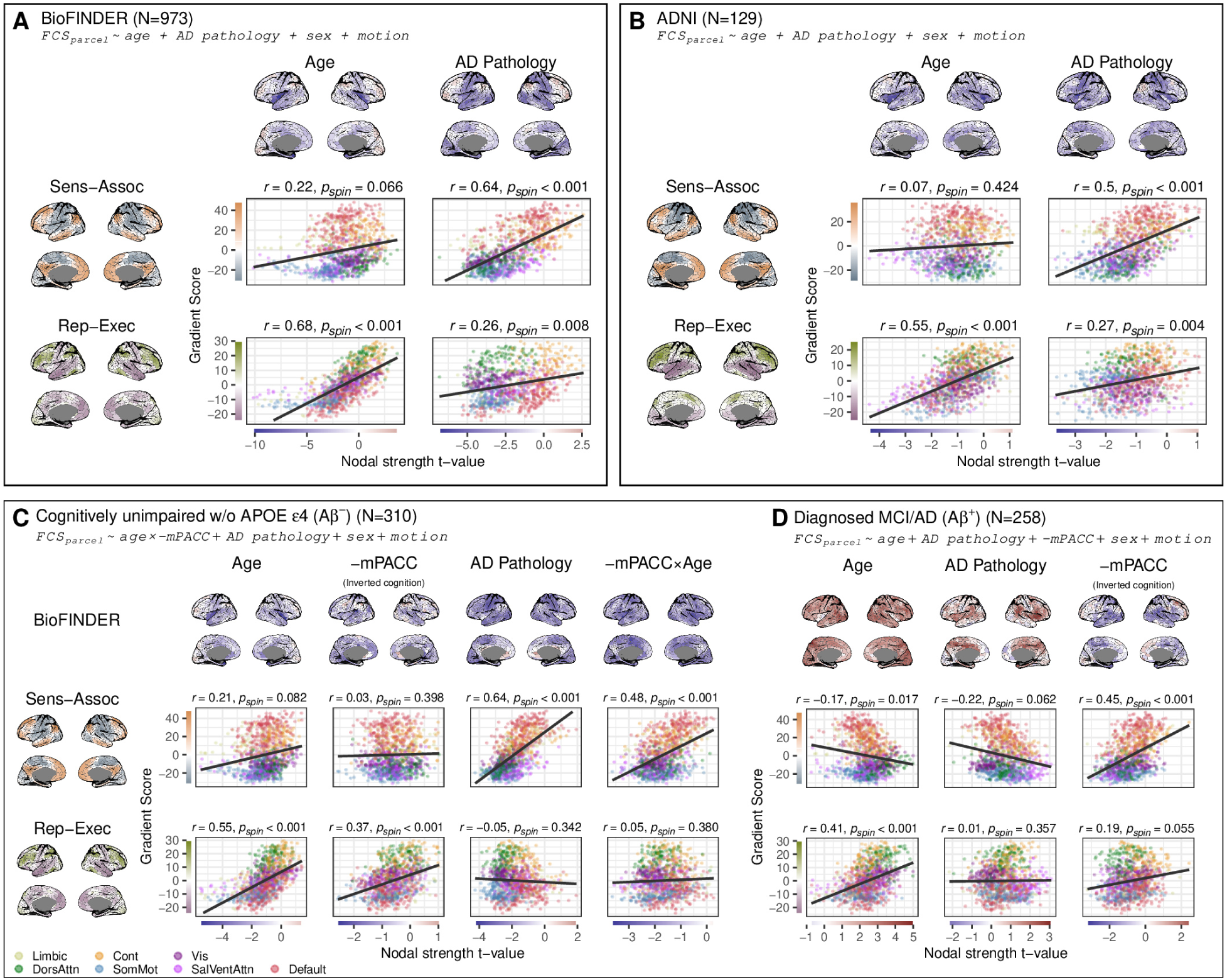
Gradient-aligned age and AD effects are replicated using nodal connectivity strength. Analyses using nodal connectivity strength (column-wise average of the unthresholded connectivity matrix) as the outcome measure show that age-related FC effects align with the RE axis, while AD pathology-related effects align with the SA axis, consistent with the main results based on nodal affinity. Results are shown for BioFINDER (A) and ADNI (B), with cognitive status effects illustrated in (C) and (D). Cortical maps display t-values from nodal linear models, and scatter plots show relationships between t-values and gradient scores, colored by network membership. Associations were quantified using Pearson correlation, with significance assessed using a spin test (one-sided, uncorrected; see Methods).

**Figure E7:**
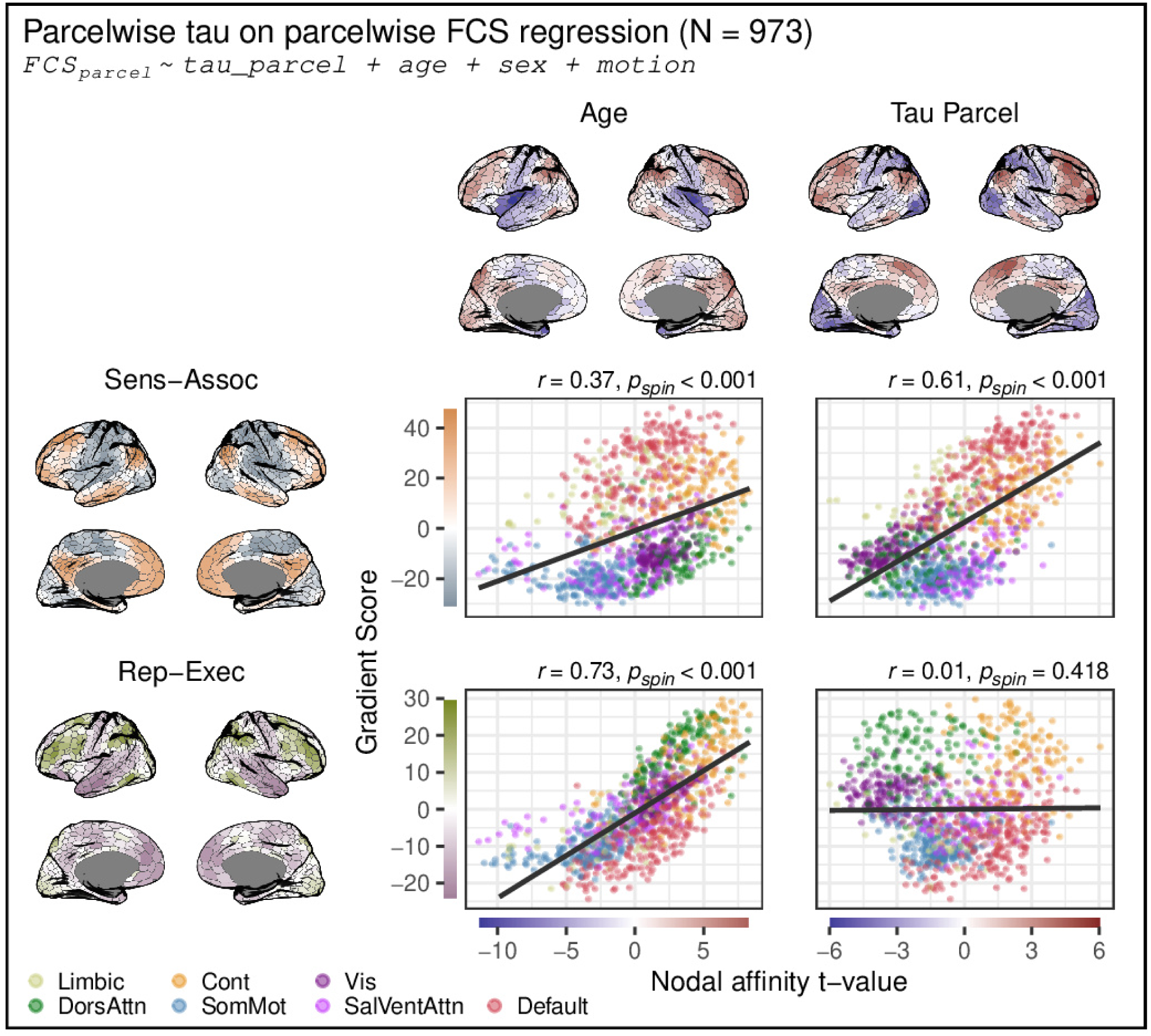
Parcel-level tau effects track global pathology patterns. Results from parcel-wise models using tau PET SUVR extracted for each Schaefer-1000 parcel (FCS_parcel_ ∼ age + tau_parcel_ + covariates). The spatial pattern closely matches analyses with global pathology measures, indicating that parcel-level tau largely tracks global tau burden on the effect of FCS.

Because nodal affinity and the SA axis share some broad structure inherent to large correlation matrices, we examined whether this could explain the pathology–gradient alignment. After removing parcel-wise variance shared with nodal affinity averaged over subjects (Figure E8 B), the alignment between pathology and the SA axis FCS persisted, indicating that it is not driven by circular dependencies in the input data. Moreover, predictors differ in their gradient alignment: age aligns with the RE axis, education shows an inverse RE alignment, and some factors (e.g., sex) show no alignment with either axis, indicating that the SA axis is not a generic attractor (see Figure E8 A for sex and education effects).

**Figure E8:**
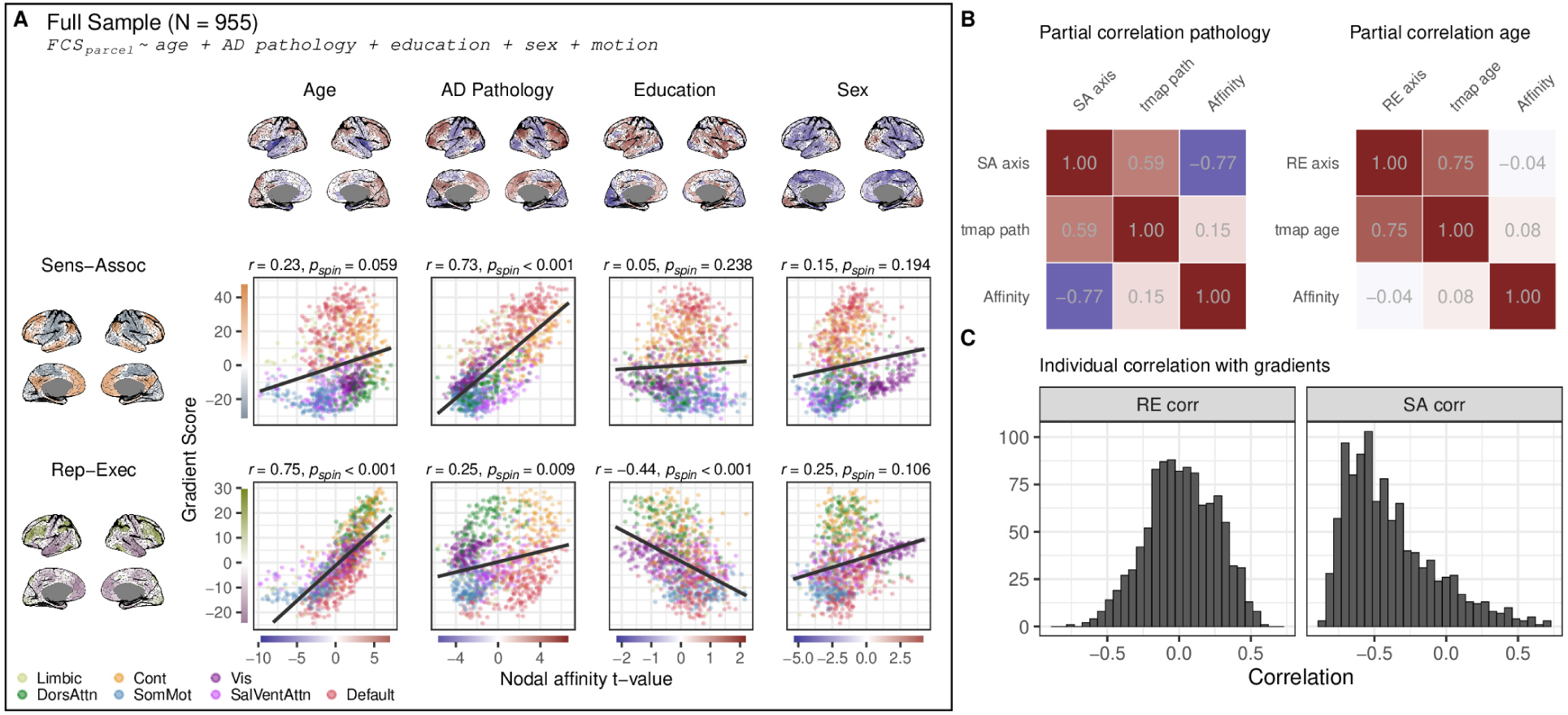
Gradient alignment of pathology persists beyond shared structure and differs across predictors. Cross-sectional results of the full sample with education in the model and sex as variable of interest (A). Correlations of the t-maps from pathology (path) and age in the model FC_parcel_ ∼ age + AD pathology + sex + motion, when accounting for the variance of nodal affinity averaged across subjects. The correlation between the pathology t-map and the SA axis persists even though we account for mean nodal affinity, indicating it is not the shared data structure that accounts for the correlation between the two (B). Individual nodal affinity maps correlated with the SA and RE axis respectively show that, while individual affinity maps tend to negatively correlate with the SA axis, they do not on average tend to correlate with the RE axis.

## Discussion

This study demonstrates that age- and AD-related reorganization in functional connectivity are fundamentally shaped by macro-scale brain organization. Our findings offer a more holistic perspective of connectivity alterations free from constraints imposed by predefined region or network borders, allowing reinterpretation of prior, often contradictory, findings. We highlight that increases or decreases in functional connectivity similarity (and strength) rarely occur in isolation; instead, they coexist systematically, forming consistent spatial patterns that align with two distinct principal axes describing functional brain organization. More (global) AD pathology is associated with decreased FCS in sensorimotor areas, and increased (or less decreased) FCS in associative areas – aligning with the sensorimotor-association axis. Higher age is associated with decreased FCS in representation areas, and increased (or less decreased) FCS in executive areas – aligning with the representation-execution axis. These results were replicated in an external cohort, confirmed longitudinally, and across multiple methodological choices. Further, we demonstrate that these organization-aligned spatial patterns shift with age and with severity of AD pathology. For instance, FCS effects aligned with the SA axis are detectable early in the process of AD pathology accumulation, even before clinically detectable cognitive impairment, but diminish once clinical impairment develops. At this later stage, SA alignment more directly reflects worse cognitive performance rather than greater AD pathology, possibly signalling cognitive aspects of disease progression. A similar relationship between SA-aligned connectivity and cognitive performance appeared in clinically unimpaired older adults. Taken together, our results suggest that connectivity changes aligned with organizational axes (specifically the sensorimotor-association axes) may be associated with cognitive difficulties, even before clinical impairment becomes evident.

The cortical patterns of FCS identified in this study illustrate a balancing scale of age-related FCS increases and decreases, anchored at the centers of the SA and RE axis for AD and aging, respectively. FCS changes in one functional domain, as captured by these gradients (e.g., decreased FCS in sensorimotor areas along the SA axis), are mirrored by less or opposing changes in the “opposite” domain (e.g., increased FCS in associative regions). This dynamic is especially noteworthy in light of the cognitive domains most affected by AD and aging, which align closely with the SA and RE axis, respectively. AD primarily impairs episodic memory^44^, and the limbic, default mode, and associative networks supporting memory lie on the end of the SA axis that tended to show increased FCS with early pathology in our data. In contrast, normal aging disproportionately affects executive functioning^45,46^, supported by the dorsal attention and control networks^47^, which lie on the executive end of the RE axis, the same side that showed increased FCS with age.

Consistent with this alignment of cognitive burden and gradient architecture, our analyses with domain specific cognitive performance showed that better memory and executive performance each exhibited FCS effects aligning with both axes. Numerically, executive performance aligned more strongly with the RE axis than the SA axis, and memory aligned more strongly with the SA axis than the RE axis. Importantly, higher performance in both memory and executive domains was associated with positive gradient alignment, in contrast to global cognition, which had negative alignment. This could suggest that gradient aligned effects may support domain specific cognitive functioning. The parcel-wise mediation analyses added to this interpretation. Specifically for AD pathology, SA-aligned effects tended to be beneficial for memory. Age related RE-alignment had a similar but less clear pattern. Importantly, the indirect effects on the two cognitive domains were inversely related, such that effects seemingly beneficial for one domain tended to be detrimental for the other. This is consistent with the notion that cognitive processes draw on shared, limited neural resources^48^. Together, these patterns point to a systematic reconfiguration of connectivity that may reflect shifts in resource allocation^2^, and highlight that macroscale gradients capture not only organizational principles of the cortex but also functional susceptibilities arising in the context of aging and AD.

Situating the findings of the present study in prior literature warrants some consideration. In our main analyses we do not assess FC strength, which is the more common FC measure in the literature, hence our results are not directly translatable to concepts such as hyper- and hypoconnectivity. However, the supplementary analysis with connectivity strength as outcome revealed that the spatial distribution of effects are very similar across measures – but with the important distinction that nodal strength showed weaker (if any) increases over the parcels, whereas the same areas using nodal affinity as outcome showed very clear increases.

With that in mind, we note that the spatial patterns of FCS effects observed here partly resemble those reported for connectivity strength, despite the considerable heterogeneity in the FC literature in aging and AD^18,40^. For instance, hypoconnectivity within the default mode^7,49^ and salience networks – particularly in the opercular insula^18^ – has been frequently reported in healthy aging. We observed analogous decreases in FCS in these same systems. Age associated increases of FC, as seen in the present study, are on the other hand not commonly reported. However, a systematic review^50^ noted increased integration across most brain systems (except primary processing), which could explain the increased FCS in executive areas presented in this study.

At a broader level of organization, our findings align with gradient-based descriptions of neurodegeneration, which reported that age and AD pathology load onto partially distinct macroscale axes of metabolism (eigenbrains)^31^. In that work, an axis resembling our RE axis, better captured aging-related variance and an axis resembling the sensory-association axis better captured AD-related variance. A similar axis-specific organization emerged in our functional connectivity results. In AD, we observed widespread pathology-related reductions in FCS in unimodal sensory–motor regions alongside increases in transmodal association cortex. This pattern closely parallels very recent findings using connectivity-strength measures^32^, providing direct convergence across modalities in showing decreased sensory connectivity and increased associative connectivity in AD. This whole-brain correspondence is highly encouraging, and echoes fragmentary evidence for the same dissociation that has been reported across prior regional connectivity studies: decreases in sensory–motor regions^51,52^ and increases in associative or DMN regions^1,10,17,19^, including reports of DMN “hyperconnectivity” in early and preclinical AD^53–55^. Notably, we did not observe consistent gradient alignment for *APOE* ε4 carriers.

Previous reports^12,17,19^ have suggested that initial hyperconnectivity early on in the disease may give way to hypoconnectivity as the disease advances. This hypoconnectivity has also been observed^17,56–58^. Parallelling this “phasic” view, our nonlinear analysis showed that gradient alignment peaked at moderate pathology levels before declining as pathology burden increased further. Our longitudinal analysis further reinforced this, demonstrating that within-subject pathology accumulation (ΔPathology) was associated with gradient-aligned FCS changes, but that this relationship disappeared for individuals with higher baseline pathology.

Strikingly, the relationship between pathology load and SA-aligned FCS was entirely absent in MCI and AD patients, a pattern that may partly reflect age–pathology interactions, whereby age-related effects modulate the expression of pathology-related SA-alignment^31^. Instead, in this group, having a lower cognitive performance was associated with an SA-like FCS pattern. A similar relationship to cognition also emerged among clinically unimpaired older adults. Previous studies have linked hyperconnectivity in the DMN with subjective cognitive decline^59^, as well as sensory-motor hyper- and hypoconnectivity with better and worse cognitive performance respectively^60,61^, again parallelling our results. Together, these findings could suggest that the SA-related FC patterns we observed represent a harmful state, indicating increased vulnerability to pathological processes leading to cognitive decline^12^. An alternative interpretation could be that individuals experiencing underlying cognitive difficulties – potentially due to co-morbidities or less cognitive resilience – require greater compensatory adaptations. These interpretations are not mutually exclusive. And both are consistent with our observation that several non-AD pathological markers were independently associated with gradient-aligned FCS changes, which suggests that these patterns are associated with broader system-level strain rather than AD pathology (or aging) alone. Intervention studies with repeated imaging and detailed cognitive testing over time^9,11,62^ will, however, be necessary to disentangle the temporal dynamics of these interactions to assess if the gradient-like FCS alterations are compensatory, harmful, or both.

Regardless of interpretation, gradient-like FCS alterations appear to signal ongoing physiological or pathological processes directly or indirectly related to cognition, as indicated by the marked SA-alignment of AD pathology (driven by tau) in the cognitively unimpaired group. Understanding how to leverage these signals clinically might enable identification of patients experiencing cognitive vulnerability before overt clinical symptoms emerge. Beyond this clinical implication, our results also offer conceptual clarity for FC research more broadly. Rather than viewing gradients as outcomes, this study demonstrates their value as interpretative frameworks for understanding how brain function adapts or responds to aging and disease processes.

The strength of this study lies in its large, single-site main cohort and the several methods employed to validate the findings within it, including an external replication, nonlinear, and longitudinal linear and nonlinear analyses. The longitudinal approach of this paper should be emphasized, as such within-subject designs are still relatively rare in the field^63^. By using a whole-brain perspective and continuous measures of AD pathology, we also overcome the limitations of region-specific analyses and categorical diagnostic classifications. Using several different methodological approaches we show that the results are generally not methodology dependent, but that differences between nodal affinity and nodal connectivity strength may be worth further investigation. However, this study also has several limitations. First, our whole-brain approach overcomes biases inherent in region- or network-specific analyses but limits the ability to pinpoint specific connections driving the gradient-aligned patterns. Second, resting-state connectivity has a weaker relationship to cognition compared to task-based paradigms^60,64^; future studies should explore whether task-based FC strengthens or refines the findings of this study. Third, as an observational study, causal inferences regarding the relationship between functional connectivity, pathology, and cognitive decline is limited. Fourth, limited cohort diversity may restrict generalisability: over 90% of BioFINDER participants were native Swedish speakers, and over 90% of ADNI participants self-identified as white. Lastly, while we stratified patients within the typical AD spectrum into amnestic and non-amnestic groups to assess differential trajectories of gradient alignment across pathology, the limited representation of atypical AD phenotypes (e.g., posterior cortical atrophy, logopenic variant primary progressive aphasia, dysexecutive AD) constrains our ability to determine whether domain-predominant cognitive impairments are associated with distinct gradient alignments or with reduced pathology-to–sensory–association alignment in advanced disease.

In sum, our findings reveal that age- and AD-related FCS alterations align with distinct large-scale cortical gradients of functional organization. These alignments vary along the age and AD pathology spectra, and are associated with cognition, independent of AD pathology burden. We propose that gradient-like FCS changes may reflect a general neural signature of cognitive strain or system-level stress, while acknowledging that the directionality of this relationship is uncertain. Similarly, we cannot conclude whether this signature is detrimental or compensatory – our parcel-wise mediation analyses seem to suggest that they are both. We show that these patterns emerge in older individuals, those that are cognitively healthy but have subtle AD pathology, and clinically impaired patients alike, suggesting that cognitive demands may increasingly exceed available neural resources in these groups.

## Acknowledgements

We would like to thank all research participants and their family members, and everyone involved in the collection, refinement, and administration of the data used in this study. Their efforts and contributions have made this research possible. JR and JW were supported by the SciLifeLab Wallenberg Data Driven Life Science Program (grant: KAW 2020.0239), the Swedish Alzheimer Foundation (AF-994626), and the Swedish Research Council (2024-03642). JR was additionally supported by the Eva and Oscar Ahrens Foundation and Strategic Research Area MultiPark at Lund University through travel grants to present this work. The BioFINDER-2 study was funded by the European Research Council (ADG-101096455), the Swedish Research Council (2022-00775, 2021-02219), Alzheimer’s Association (ZEN24-1069572, SG-23-1061717), GHR Foundation, ERA PerMed (ERAPERMED2021-184), Knut and Alice Wallenberg foundation (2022-0231), Strategic Research Area MultiPark (Multidisciplinary Research in Parkinson’s disease) at Lund University, Swedish Brain Foundation (FO2021-0293, FO2023-0163), Parkinson foundation of Sweden (1412/22), WASP and DDLS Joint call for research projects (WASP/DDLS22-066), Swedish Alzheimer Foundation (AF-1011799, AF-980907, AF-994229, AF-1032702), Cure Alzheimer’s fund, Rönström Family Foundation, Berg Family Foundation, Konung Gustaf V:s och Drottning Victorias Frimurarestiftelse, Skåne University Hospital Foundation (2020-O000028), Regionalt Forskningsstöd (2022-1259) and Swedish federal government under the ALF agreement (2022-Projekt0080, 2022-Projekt0107). The precursor of 18F-flutemetamol was sponsored by GE Healthcare. The precursor of 18F-RO948 was provided by Roche. TDS was supported by grants from the National Institute of Health (R37MH125829, R01MH120482, R01EB022573, R01MH112847, U24NS130411), the Penn-CHOP Lifespan Brain Institute, and the AI2D Center. NF was supported by the BrightFocus Foundation (A2021026S), the Gerhard and Ilse Schick Foundation and the Alzheimer’s Forschung Initiative (23074CB). LC was supported by Fondation Philippe Chatrier. LEMW was supported by the Bente Rexed Gerstedt Foundation, the Hains Foundation and MultiPark, a strategic research area at Lund University. HHB was supported by the European Union’s Horizon Europe Research and Innovation Program under Marie Sklodowska-Curie action (101153323). The funding sources had no role in the design and conduct of the study; in the collection, analysis, interpretation of the data; or in the preparation, review, or approval of the manuscript.

Data collection and sharing for the Alzheimer’s Disease Neuroimaging Initiative (ADNI) is funded by the National Institute on Aging (National Institutes of Health Grant U19AG024904). The grantee organization is the Northern California Institute for Research and Education. In the past, ADNI has also received funding from the National Institute of Biomedical Imaging and Bio-engineering, the Canadian Institutes of Health Research, and private sector contributions through the Foundation for the National Institutes of Health (FNIH) including generous contributions from the following: AbbVie, Alzheimer’s Association; Alzheimer’s Drug Discovery Foundation; Araclon Biotech; BioClinica, Inc.; Biogen; Bristol-Myers Squibb Company; CereSpir, Inc.; Cogstate; Eisai Inc.; Elan Pharmaceuticals, Inc.; Eli Lilly and Company; EuroImmun; F. Hoffmann-La Roche Ltd and its affiliated company Genentech, Inc.; Fujirebio; GE Healthcare; IXICO Ltd.; Janssen Alzheimer Immunotherapy Research & Development, LLC.; Johnson & Johnson Pharmaceutical Research & Development LLC.; Lumosity; Lundbeck; Merck & Co., Inc.; Meso Scale Diagnostics, LLC.; NeuroRx Research; Neurotrack Technologies; Novartis Pharmaceuticals Corporation; Pfizer Inc.; Piramal Imaging; Servier; Takeda Pharmaceutical Company; and Transition Therapeutics.

## Author contributions

JR and JWV conceptualized the study. JR, LC, HHB, LEW, OH and JWV contributed to analytical design. JR, NF, OS, AD, DVW, SJ, ES were involved in data processing and curation. SP, ES, RO, NMC and OH acquired and provided the data. JR conducted data analysis. JWV supervised the study. JR and JWV wrote the manuscript. All authors interpreted the data and contributed substantively to revising the manuscript

## Methods

### Sample

#### BioFINDER

Information regarding recruitment, diagnostic criteria, and Aβ positivity assessment for the BioFINDER-2 cohort (https://biofinder.se/two; clinical trial number NCT03174938) has been described in detail previously^65^. All participants provided written informed consent in accordance with the Declaration of Helsinki. Ethical approval for the study was obtained from the Ethics Committee of Lund University, Lund, Sweden, with additional permissions for PET imaging granted by the Swedish Medicines and Products Agency and the local Radiation Safety Committee at Skåne University Hospital, Sweden. Participants in BioFINDER-2 are compensated with 400 SEK per visit/examination (MRI, PET, LP) to cover travel costs to and from the examinations. For the cross-sectional analysis, 1,168 individuals had all relevant data available (fMRI, CSF Aβ42/40, and tau PET). However, as described in the Imaging section, a stringent in-scanner motion filter was applied, excluding 194 individuals and leaving a final cross-sectional sample of N = 973. Of these, 655 were cognitively unimpaired or had self-reported cognitive decline (SCD), while 319 were diagnosed with mild cognitive impairment or dementia^66^. Cognitively impaired cases were included only if they were Aβ-positive, as assessed by abnormal CSF Aβ42/40 values, see the bivariate distribution of age and pathology score of these individuals in Figure E9. For the longitudinal analysis, 480 individuals had fMRI and tau PET data from at least two visits. After applying the motion filter, 378 unique individuals with at least two data points remained. See tables Table 1 and Table 2 for additional details.

**Figure E9:**
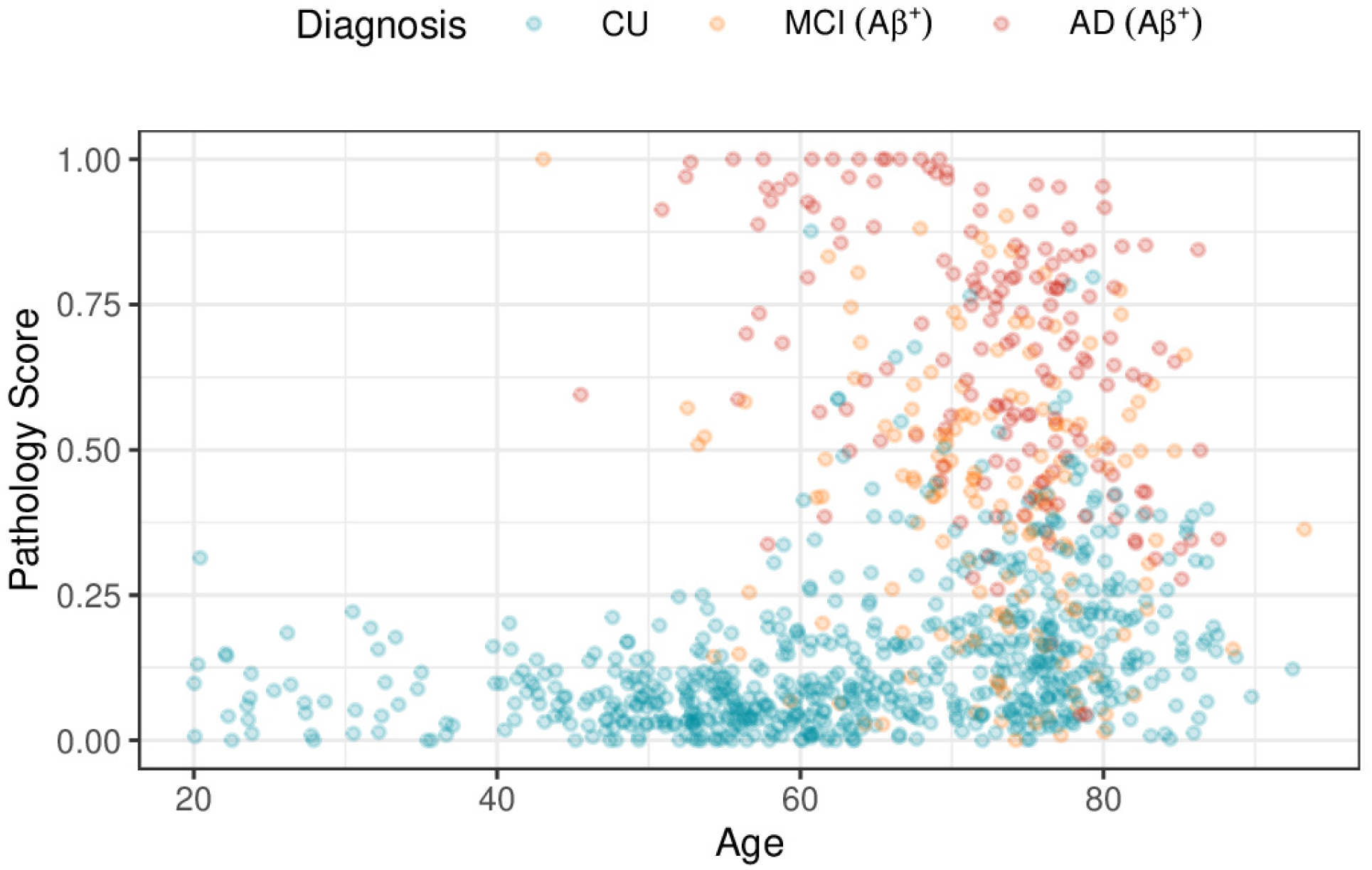
Distribution of age and pathology score of the main cross-sectional sample.

#### ADNI

Data for the validation cohort were obtained from the Alzheimer’s Disease Neuroimaging Initiative (ADNI) database (adni.loni.usc.edu). The ADNI was launched in 2003 as a public-private partnership, led by Principal Investigator Michael W. Weiner, MD, with the goal of testing whether serial MRI, PET, other biological markers, and clinical and neuropsychological assessments can be combined to measure the progression of MCI and early AD. The current goals include validating biomarkers for clinical trials, improving generalizability of ADNI data, and providing data on the diagnosis and progression of Alzheimer’s disease to the scientific community. For up-to-date information, see adni.loni.usc.edu. A total sample of N = 131 had the exact same variables available as in the discovery cohort (CSF Aβ42/40, tau PET, fMRI), with two individuals being excluded for excessive in-scanner motion. Of those, there were 89 healthy controls and 40 participants diagnosed with dementia or mild cognitive impairment. All cognitively impaired cases were Aβ-positive as assessed by abnormal CSF Aβ42/40 or amyloid PET. See table Table 1 for further details.

## Data acquisition and processing

### CSF and *APOE* ε4 carriership

#### BioFINDER

CSF Aβ42 and Aβ40 levels in BioFINDER were measured using the Roche Elecsys platform (Roche Diagnostics International Ltd., Basel, Switzerland). Aβ positivity was determined by Gaussian mixture modeling, using a cutoff of <0.080, previously described in^65^. Participants who carried at least one copy of the *APOE* ε4 allele were defined as *APOE* ε4 carriers.

#### ADNI

CSF Aβ42 and Aβ40 levels in ADNI were measured using the Roche Elecsys platform with Elecsys CSF immunoassays on a cobase 601 analyzer at the University of Pennsylvania, as described in^67^. Aβ positivity was determined using previously established amyloid-PET cut-offs derived in^39^. *APOE* genotyping was performed using blood samples, and participants carrying at least one *APOE* ε4 allele were classified as *APOE* ε4 carriers. See^67^ for further details on CSF analysis and APOE genotyping in ADNI.

### Cognitive assessment

Estimation of cognitive performance using the modified Preclinical Alzheimer’s Cognitive Composite (mPACC^42^) in BioFINDER has been previously described^66^. It is the average of five z-scores from the cognitive tests: Alzheimer Disease Assessment Scale (ADAS) delayed recall (weighted double), Animal Fluency, Mini-Mental State Examination (MMSE) and Symbol Digit Modalities Test (SDMT).

To derive cognitive domain scores while accounting for shared variance across tests, we estimated McDonald’s omega using a bifactor model with Schmid–Leiman transformation using the omega() function in the R package psych. Ten cognitive tests were included: ADAS immediate and delayed recall, Trail Making Test A and B, Symbol Digit Modalities Test, AQT Color–Form, VOSP Cube, Animal Fluency, Letter Fluency, and the Boston Naming Test. A four-domain structure (memory, executive, visuospatial, and language) was specified to capture cognitive patterns observed in both typical Alzheimer’s disease and atypical presentations, including posterior cortical atrophy (predominant visuospatial impairment), logopenic variant primary progressive aphasia (language impairment), and dysexecutive Alzheimer’s disease (executive dysfunction). The model was estimated on the test correlation matrix (Pearson) from cognitively healthy participants under 70 years (n=285) using minimum residual (OLS) factor extraction, with an oblique (oblimin) rotation applied prior to Schmid–Leiman transformation. This yields a general cognitive factor (g) and domain factors defined by the specified structure. Regression-based factor scores for g and each domain, for each individual in the full sample, were extracted from the Schmid–Leiman solution. Component loadings can be seen in Figure 6. Out of the original 973 participants 118 had missing data on one or more test scores, 91 of these were AD or MCI. Individuals with more than 5 test scores missing were filtered out. The remaining missing values were handled using multivariate imputation by chained equations (MICE). Age, sex, education, diagnosis, AD pathology, and mPACC were included as predictors. Each test was imputed using the remaining tests together with these covariates, using predictive mean matching. Twenty imputations with 30 iterations per chain were run, and convergence was assessed using trace plots. A single complete dataset was obtained by averaging the imputed values across imputations.

An individual was defined to have a domain impairment if their domain score was below one standard deviation of the mean of the healthy controls. If a subject had an impairment on the memory domain they were classified as amnestic. If a subject did not have an impairment on memory but on any other domain they were classified as non-amnestic.

### Pathology score

As with our previous work^68^, a pathology score representing accumulated AD neuropathology was calculated using the R package SCORPIUS^37^. The variables included were the CSF Aβ42/40-ratio, together with PET SUVR from regions reflecting Braak I-II, III-IV and V-IV as defined by^69^. SCORPIUS is a trajectory inference algorithm designed to nonlinearly project high-dimensional data onto a single continuous path. Briefly, pairwise Euclidean distances between observations were calculated. The resultant distance matrix was then decomposed and three components retained. Each individual is hence represented by coordinates in this three-dimensional space. Observations were then clustered into four clusters using k-means and the shortest path going through all cluster centers found. A principal curve was then iteratively fitted to the data to model the progression trajectory. Methodological parameters, such as the number of compo-nents and cluster centers, were set to the default values used in SCORPIUS. Each participant was projected onto this curve to assign a pathology score reflecting its position along the inferred trajectory. See Figure 1 D for estimation of the pathology measures as a function of the combined pathology score and distribution of patient groups over it. For the longitudinal pathology score, since longitudinal CSF Aβ40/42 data was largely unavailable, only tau PET SUVR from the three Braak meta ROIs (I-II, III-IV and V-VI) were used. The pathology scores were derived separately within each cohort.

## Imaging

### MRI and Tau PET

#### BioFINDER

Structural MRI was acquired using a Siemens 3T MAGNETOM Prisma scanner (Siemens Healthineers, Erlangen, Germany) with a 64-channel head coil. T1-weighted images (Magnetization Prepared – Rapid Gradient Echo, MPRAGE) were acquired with the following parameters: in-plane resolution = 1 × 1 mm², slice thickness = 1 mm, repetition time (TR) = 1900 ms, echo time (TE) = 2.54 ms, flip-angle = 9°.

All resting-state fMRI data (eyes closed) were acquired using a 3D echo-planar imaging (EPI) sequence with an in-plane resolution of 3×3 mm and slice thickness 3.6 mm; echo time = 30ms, and flip-angle = 63°. Scan time was 7.85 minutes, with a multiband repetition time of 1020 ms, resulting in 462 frames per scan before processing and censoring. Preprocessing has briefly been described in^15^. The processing was performed using a modified CPAC^70^ pipeline, building mostly on FSL^71^, AFNI^72^ and ANTS^73^. Skullstripping was done with in-house code using the T2 structural image as a primer and Brain Surface Extractor from BrainSuite^74^. The fMRI preprocessing included slice-timing and motion correction. Physiological noise was regressed out using CompCor^75^, alongside the removal of linear and quadratic trends. Additionally, regression of Friston’s 24-parameter motion correction^76^, white matter and CSF signals were performed. Susceptibility distortion was corrected by unwarping the functional data using a nonlinear diffeomorphic transformation (performed with ANTS) of the mean functional image to high resolution (1 × 1 × 1 mm3) T2 structural image^77^. This transformation was then applied to each volume in the fMRI timeseries. Finally, a bandpass filter (0.01-0.1 Hz) was applied, and images were transformed from native to MNI space with ANTS. For each scan, frames were censored based on DVARS for that frame lying outside 1.5*IQR above and below the third and first quartiles, respectively^78^. To avoid distortion from the outlier frames, they were interpolated before bandpass filtering and then removed from the final 4D image. Finally, any participant with an average FD > 0.3mm or maximum FD > 3mm over the entire sequence were filtered out before analysis.

Tau PET acquisition has been described in detail by^79^. Briefly, participants were injected with an average of 365 ± 20 MBq [18F]RO948, and emission data was acquired between 70 and 90 minutes post-injection, adjusted for the tracer’s pharmacokinetics. Low-dose CT scans were conducted before PET scans for attenuation correction. PET data was reconstructed using the VPFX-S algorithm (ordered subset expectation maximization with time-of-flight and point spread function corrections). The LIST mode data was binned into 4 × 5-minute time frames, and images were motion-corrected, summed, and co-registered to corresponding T1-weighted MRI images. FreeSurfer parcellation (v6; http://surfer.nmr.mgh.harvard.edu.ludwig.lub.lu.se/) was applied to extract regional uptake values, normalized to the mean uptake in inferior cerebellar grey matter.

#### ADNI

Image processing for ADNI has been described in^80^. Briefly, MRI data were collected using 3T scanners with unified scanning protocols that can be accessed here: https://adni.loni.usc.edu/wp-content/uploads/2017/07/ADNI3-MRI-protocols.pdf. Structural MRI was acquired with a 3D T1-weighted MPRAGE sequence (1 mm isotropic voxels, TR = 2300 ms).

Two protocols exist for the resting state EPI-BOLD sequence in ADNI: Basic and Advanced. De-pending on which protocol a participant has been subjected to, the parameters vary. Both protocols have an approximate scanner time of 10 minutes. The Basic protocol has a TR/TE/flip angle=3000/30/90° and the Advanced a TR/TE/flip angle=600/30/53°, so the number of frames differ depending on the protocol and ranges from approximately 200 to 1000. Processing steps described in^80^ included motion correction, as well as regression of six motion parameters and mean signal from CSF and white matter. Trends were removed and a (0.01–0.08Hz) band-pass filter was applied. Images were nonlinearly registered to MNI space with coregistration to the baseline T1 image. After timeseries had been extracted, rows from high motion frames (> 0.5mm framewise displacement) were censored. These rows, together with one preceding and two subsequent, were removed. Out of the 131 subjects with a full set of variables (CSF Aβ 40/42, tau PET, fMRI), one individual was filtered out due to a maximum number of frames < 100 and one participant with an average FD > 0.3mm or maximum FD > 3mm over the entire sequence was filtered out before analysis.

Tau PET imaging in ADNI3 was performed using [^18F]AV-1451 (Flortaucipir). Participants received an injection of 370 MBq (±10%), and emission data was acquired 75 minutes post-injection. Scans were conducted for 30 minutes in six 5-minute frames. Images were reconstructed using iterative algorithms specific to each scanner model, incorporating corrections for scatter, attenuation, and motion artifacts. CT scans or transmission scans were performed for attenuation correction prior to emission imaging. More detailed information can be accessed here https://adni.loni.usc.edu/wp-content/uploads/2012/10/ADNI3_PET-Tech-Manual_V2.0_20161206.pdf.

### Assessment of functional connectivity (similarity)

In both BioFINDER and ADNI, parcel-wise timeseries were extracted using the 1000-region Schaefer atlas^33^, a cortical parcellation based on functional connectivity patterns and spatial contiguity. The timeseries were smoothed using a full-width at half-maximum kernel of 6 mm in BioFINDER and 4 mm in ADNI. After mean centering and scaling the raw timeseries, pairwise Pearson correlations were computed between parcels, resulting in 1000**×**1000 correlation matrices representing co-activation patterns. Negative correlations were set to zero. FC was quantified primarily as nodal affinity^20^, where each connectome was column-wise thresholded to retain the 25% strongest connections, and cosine similarity was calculated between columns to yield a matrix of connectivity similarity. This approach shifts the interpretation from similarity of brain activity to similarity of connectivity patterns. Margulies et al.^20^ used a threshold retaining the 10% strongest connections, while we opted for a more liberal threshold due to the lower spatial resolution in our clinical data and coarser brain parcellation of using 1000 parcels instead of surface vertices.

Nodal affinity reduces the rank of the original correlation matrix^81^, highlighting dominant patterns of variance while preserving interpretability. Averaging the affinity matrix across parcels then yields a metric that reflects how generic or specialized a parcel’s connectivity profile is relative to the rest of the brain: higher values indicate widespread similarity, whereas lower values point to more distinct, unique connectivity patterns. Compared to nodal strength, the parcel-wise average of correlation coefficients, affinity emphasizes the configuration of connections rather than their overall magnitude. This makes it a particularly interesting measure in the context of aging and AD, where hypotheses often focus on loss of differentiation or “integration” of brain regions^50^.

Affinity is also widely used in functional gradient studies because it is traditionally the underlying matrix from which the gradients are derived^20^. For comparison, nodal connectivity strength – the parcel-wise average of the original connectivity matrices – was also calculated. All main analyses were replicated with nodal strength as the outcome, yielding results comparable to those with nodal affinity (see Figure E6). Additionally, we replicated our findings using correlation instead of cosine similarity on unthresholded connectivity matrices, which also yielded comparable results (see Figure S12). In Figure S10 and Figure S11, we run the main analyses with nodal affinity calculated within and between each Yeo 7 network, respectively, by taking the average affinity values of each parcel within its own network, and the average of each parcel’s affinity to parcels outside of its own network.

### Derivation of functional gradients

Breaking down the connectivity matrices into low-dimensional components, known as gradients^20^, captures connectivity patterns of the brain as continuous measures of connectivity similarity over the cortex. Regions with similar connectivity patterns will get grouped together at different ends of these gradients. The primary components in such analyses represent the major variance axes of functional connectivity. For BioFINDER and ADNI, respectively, we calculated an estimate of the functional connectome by averaging individual connectivity matrices from cognitively unimpaired *APOE* ε4 non-carriers without abnormal Aβ levels. In BioFINDER, only younger participants (≤60 years) were included (n = 125), whereas in ADNI, the entire age range was used (n = 82) due to the unavailability of a younger sample.

The variance components, or gradients, of these two connectomes were derived using principal component analysis (PCA) with singular value decomposition directly on the connectome without thresholding. Traditionally, gradients are derived using diffusion map embedding^20,82^, which involves applying heavy thresholding to the connectome before taking the pairwise cosine similarity of the rows/columns. However, we found that both methods produced comparable components with respect to the original gradients from^20^ (accessed from https://identifiers.org/neurovault.collection:1598). We felt that applying PCA directly on the connectome, without transformation or thresholding, offered a simpler and more interpretable approach compared to diffusion map embedding, and did not result in a notable divergence. In Figure E10, we compare the two methods over a set of thresholding values, supporting this decision (see also^83^ for a more formal comparison). Supplementary sensitivity analyses using gradients derived from both diffusion map embedding and PCA over different thresholds confirmed that this choice did not alter our results to any great extent, see Table S1.

**Figure E10:**
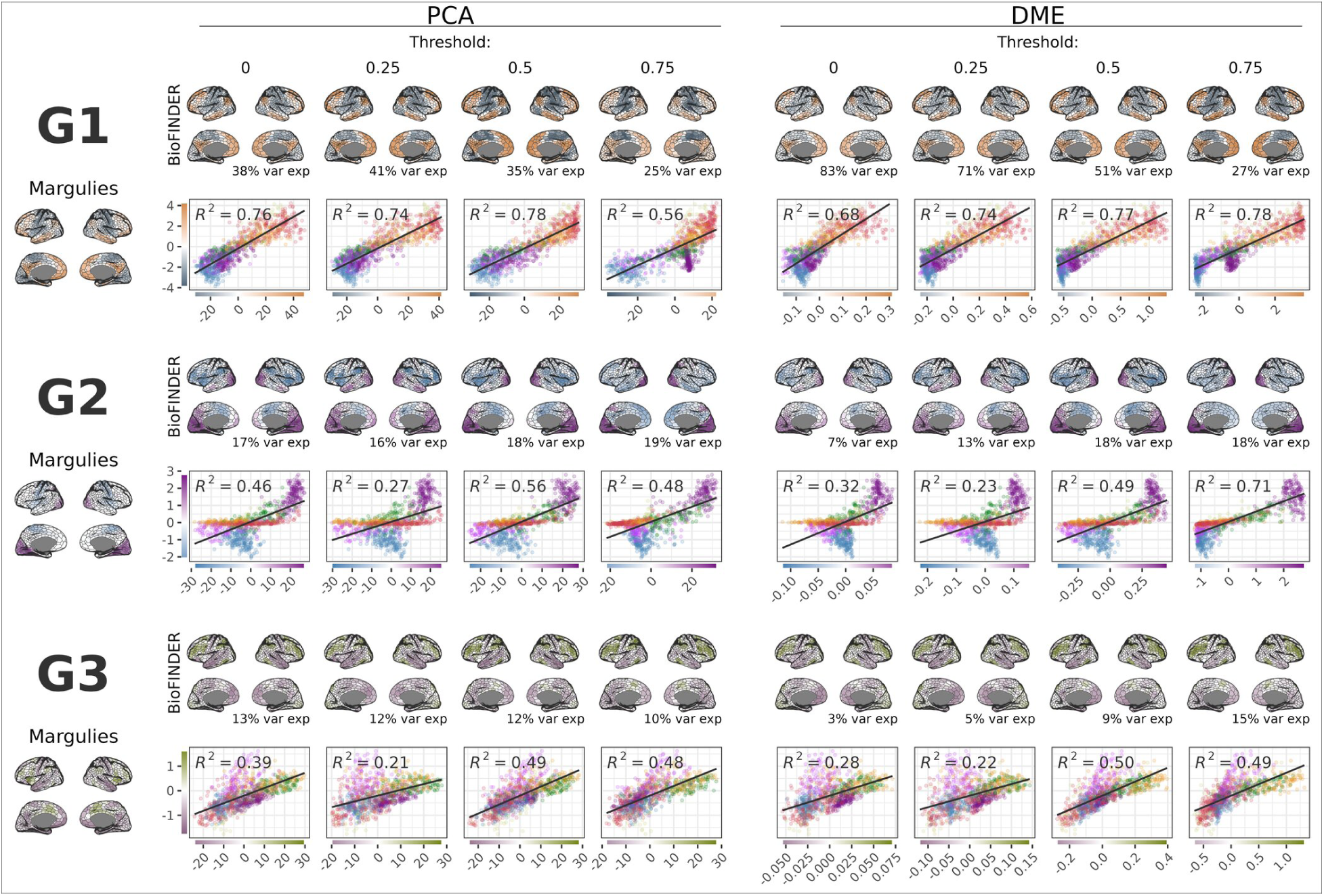
Derived gradients compared over a set of different parameter values show that the derivation method (PCA or DME) only has marginal effect on the overall spatial structure of the gradients. Reference gradients from^20^ are presented in the leftmost column. PCA method: principal component analysis via singular value decomposition on the average connectivity matrix from individuals BioFINDER without cognitive impairment, *APOE* ε4 non-carriers without abnormal amyloid levels, and ≤60 years of age. DME: diffusion map embedding (DME). The gradients using either method are shown for 4 different levels of FC thresholding.

Meta-analytic decoding was performed using the NeuroQuery database^35^ to identify cognitive terms associated with each Schaefer-1000 parcel. For each parcel, regional associations with NeuroQuery terms were estimated using the ROIAssociationDecoder in NiMARE. We then reorganised these parcel-level results into term-level summaries by pooling, for each term, all of its parcel associations. The resulting term set was filtered to include only cognitive concepts listed in the Cognitive Atlas^36^, matched by approximate string similarity, and visualized as a word cloud highlighting their spatial alignment along the principal functional gradient in Figure E1. We first kept only words with r > 0.05. For each word, we counted how many parcels it was associated with and computed its mean gradient value. Words were assigned to a gradient side (e.g., sensory or association depending on the gradient of interest) based on this value, and within each side the 25 most frequently “mentioned” words (i.e., the words associated with the largest number of parcels) were retained.

### Statistical analyses

For the cross-sectional analyses, linear regression models were run for each parcel, with the functional connectivity metric as outcome, and the terms of interest as predictors, covaried for sex and motion (framewise displacement). For example, for the analyses looking at age and AD pathology, the model formula was as follows: *FC_parcel_* ∼ *age* + *pathology* + *sex* + *motion*. This was repeated for each of the 1000 parcels. The t-maps were then compared to the gradients using Pearson correlation. To assess the spatial correspondence between these two maps, while accounting for spatial autocorrelation, we performed a spin test based on the method introduced by^34^. Using the Hungarian algorithm, we generated 1000 random rotations of the parcel centroid coordinates, separately rotating the left and right hemispheres while preserving their anatomical structure. For each rotation, parcels were reassigned by minimizing the Euclidean distance between original and rotated centroids. We then recomputed the correlation between the rotated maps and compared the empirical correlation against the null distribution to obtain a permutation-based p-value. This p-value is defined as the proportion of permuted correlations more extreme than the empirical value in the direction of the observed effect. As the test is based on a finite number of rotations, p-values are discrete (minimum *1*/ (*N_perm_* + *1*)) and should be interpreted as permutation-based estimates. No correction for multiple comparisons was applied; instead, results were interpreted based on effect size and consistency across analyses.

To investigate cross-sectional nonlinearities in the relationship between functional connectivity and age or pathology, generalized additive models (GAMs) with penalized regression splines (thin plate) were fitted for each parcel, with age and pathology as smooth terms and sex and motion as linear covariates (*FC_parcel_* ∼ *s* (*age*) + *s* (*pathology*) + *sex* + *motion*). The models were fitted using the R package mgcv (version 1.9-1), with default values for the number k basis functions. The first derivatives of the smooth terms were calculated at 100 evenly distributed points along each spectrum using finite differences, capturing the slope and effect direction at specific ages and pathology loads. For age, these derivatives were calculated over the central 90% of the range. These derivatives were then used to calculate R² values by correlating the slopes at each point with the SA axis (for pathology) or the RE axis (for age) scores. Gradient alignment could then be assessed continuously across the age and pathology spectra. To visualize these effects, derivatives were averaged over quartiles of pathology and age. The resulting average slopes were mapped onto the cortical surface. Additionally, FC metric predictions were generated across the same ranges, holding age constant at the sample mean (66.5) when assessing pathology effects and pathology constant at 0.1 when assessing age effects. Parcels were grouped into 20 equally sized bins (ventiles) based on the SA axis (for pathology) or the RE axis (for age) scores, and predicted values within each ventile were averaged.

For the longitudinal analysis, linear mixed-effects models were fit at each parcel with FCS as the outcome and baseline age, baseline pathology, and within-subject pathology change (Δpathology) as fixed effects. Time between baseline and follow-up scan, sex and scanner motion were included as (fixed effects) covariates of no interest, and random intercepts were specified for subjects, i.e. *FC_parcel_* ∼ *age_baseline_* + *pathology_baseline_* + *Δpathology_ti_*_−_*_t0_* + *time* + *sex* + *motion* + (*1*|*subject*). Variance inflation factors in this model were all <1.21. The within-subject change in pathology was calculated as the difference between the baseline pathology score and the pathology score at each follow-up time point. Δpathology therefore captures absolute within-person change, enabling us to test whether increases in pathology are associated with concurrent changes in FCS. This approach separates the between and within subject effects. Time was included to account for unspecific longitudinal effects and differences in follow-up intervals, ensuring that the Δ pathology term captures pathology-specific within-person variation. Obtained parcel-wise t-values from each fixed effect of interest were then correlated with the parcel-wise component scores of the gradients.

To assess nonlinear patterns of longitudinal change across the old age and AD pathology spectrum, we employed a sliding window approach. This allowed us to detect localized changes without smoothing over transient effects. In this approach the model described in the previous paragraph was fitted at the parcel level repeatedly in windows traversing over baseline age and baseline pathology separately (i.e. age windows were only filtered on age and vice versa). Age windows were set at 25 years and incremented by 1 year, while pathology windows spanned 0.35 pathology score units and were incremented by 0.015, resulting in 44 and 45 windows respectively. Although relatively arbitrary, the window size of age was chosen to balance the number of subjects in each window, being sufficiently wide to capture the age effects we observed across the whole sample while being sufficiently narrow to capture nonlinearities across it. This window size turned out to be approximately 35% of the full age range, hence the pathology window was set at 35% of the pathology range [0, 1] as well. The pathology increment was chosen to get approximately the same number of windows as for age. Obtained t-values for each fixed effect of interest, in each window, were correlated with gradient scores. This allowed for a dynamic assessment of gradient alignment across both the baseline age and pathology spectra.

To examine whether FCS mediated the relationships between age or AD pathology and domain specific cognitive performance, we conducted parcel-wise mediation analyses across the cortical surface. For each cortical parcel, we specified a mediation model in which either age or AD pathology served as the independent variable (X), parcel-wise FCS as the mediator (M), and cognitive performance as the outcome (Y). Mediation models were fitted separately for memory and executive domain scores. All models included sex and mean framewise displacement as covariates, and age or pathology was additionally included as a covariate when not serving as the independent variable. For each parcel, we estimated the direct effect (X → Y), the mediator effect (X → M), and the indirect effect (X → M → Y). Indirect effects were summarized as parcel-wise mediation coefficients and mapped across the cortex (see Figure 6).

## Data availability

Data from the validation cohort ADNI is a public access dataset and can be obtained by application from http://adni.loni.usc.edu/. The BioFINDER data are not publicly available, but anonymized data may be made available upon request to qualified academic investigators. Data sharing must comply with EU General Data Protection Regulation (GDPR), decisions of the Swedish Ethical Review Authority, and regulations of Region Skåne.

## Code availability and used software

Timeseries extraction from fMRI Nifti images was performed using Nilearn and meta-analytic decoding was performed using NiMARE (0.6.0) in Python 3.12, all other analyses and visualisation was done using R 4.4.2 (2024) using an array of different packages including: boot (1.3-32), broom (1.0.8), clue (0.3-66), confintr (1.0.2), conflicted (1.2.0), cowplot (1.1.3), dplyr (1.1.4), finalfit (1.0.8), flextable (0.9.7), freesurferformats (1.0.0), gganimate (1.0.11), ggforce (0.5.0), ggnewscale (0.5.1), ggplot2 (3.5.2), ggpmisc (0.6.1), ggpp (0.5.8-1), ggpubr (0.6.0), ggraph (2.2.2), ggrastr (1.0.2), ggsci (3.2.0), ggside (0.3.1), ggtext (0.1.2), ggwordcloud (0.6.2), gratia (0.10.0), gtable (0.3.6), httr2 (1.2.1), janitor (2.2.1), jsonlite (2.0.0), kableExtra (1.4.0), knitr (1.50), lme4 (1.1-37), magick (2.8.6), matrixStats (1.5.0), mediation (4.5.1), mice (3.17.0), mixtools (2.0.0.1), patchwork (1.3.0), pbapply (1.7-2), pbmcapply (1.5.1), performance (0.15.2), plotGMM (0.2.2), png (0.1-8), ppcor (1.1), proxy (0.4-27), proxyC (0.5.2), psych (2.5.6), purrr (1.0.4), quarto (1.5.1), ragg (1.4.0), RColorBrewer (1.1-3), readr (2.1.5), readxl (1.4.5), renv (1.0.7), reticulate (1.42.0), rlang (1.1.6), rmarkdown (2.29), RSpectra (0.16-2), scales (1.4.0), SCORPIUS (1.0.9), sf (1.1-0), stringdist (0.9.15), stringr (1.5.1), tibble (3.2.1), tidygraph (1.3.1), tidyr (1.3.1), tidyverse (2.0.0), withr (3.0.2), base (4.5.3), grid (4.5.3), Matrix (1.7-4), mgcv (1.9-4), parallel (4.5.3), stats (4.5.3), tools (4.5.3), utils (4.5.3). Code for all analyses and visualisation is available at https://github.com/DeMONLab-BioFINDER/fc_changes_follow_gradients.

---

**Figure S1:**
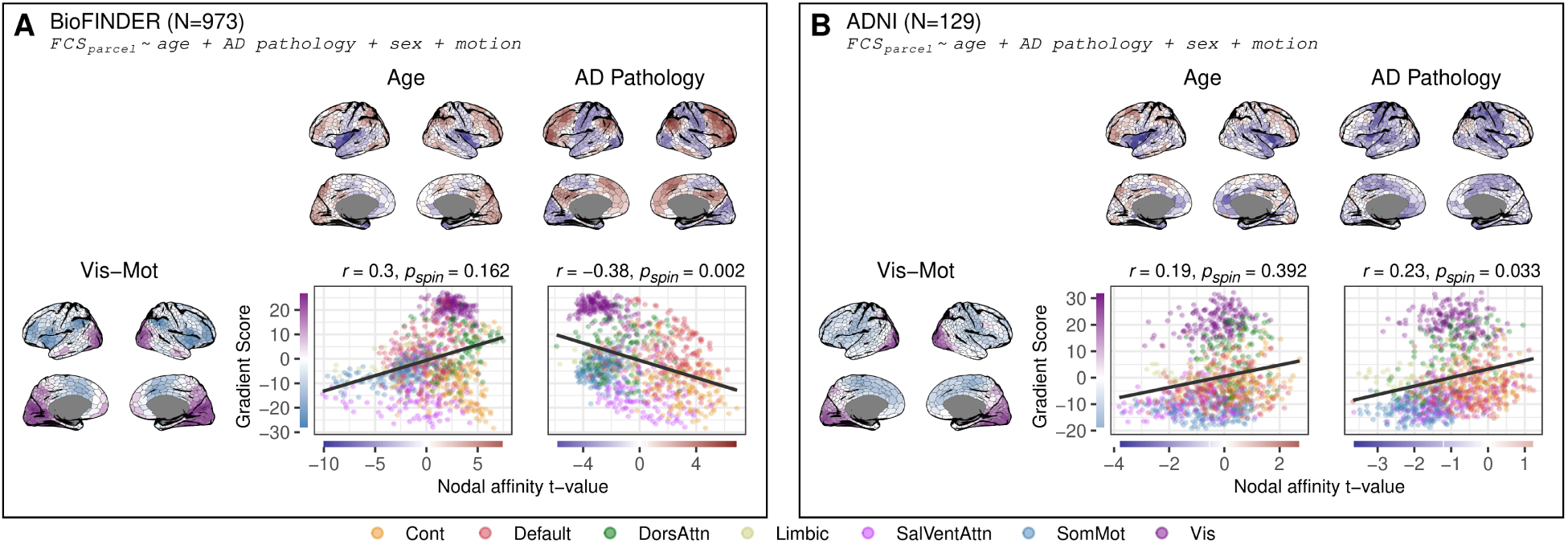
Main results of Figure 2 shown with Gradient 2. This analysis showed that there was no strong or reproducible relationships between the the t-maps for age and AD pathology and Gradient 2.

**Figure S2:**
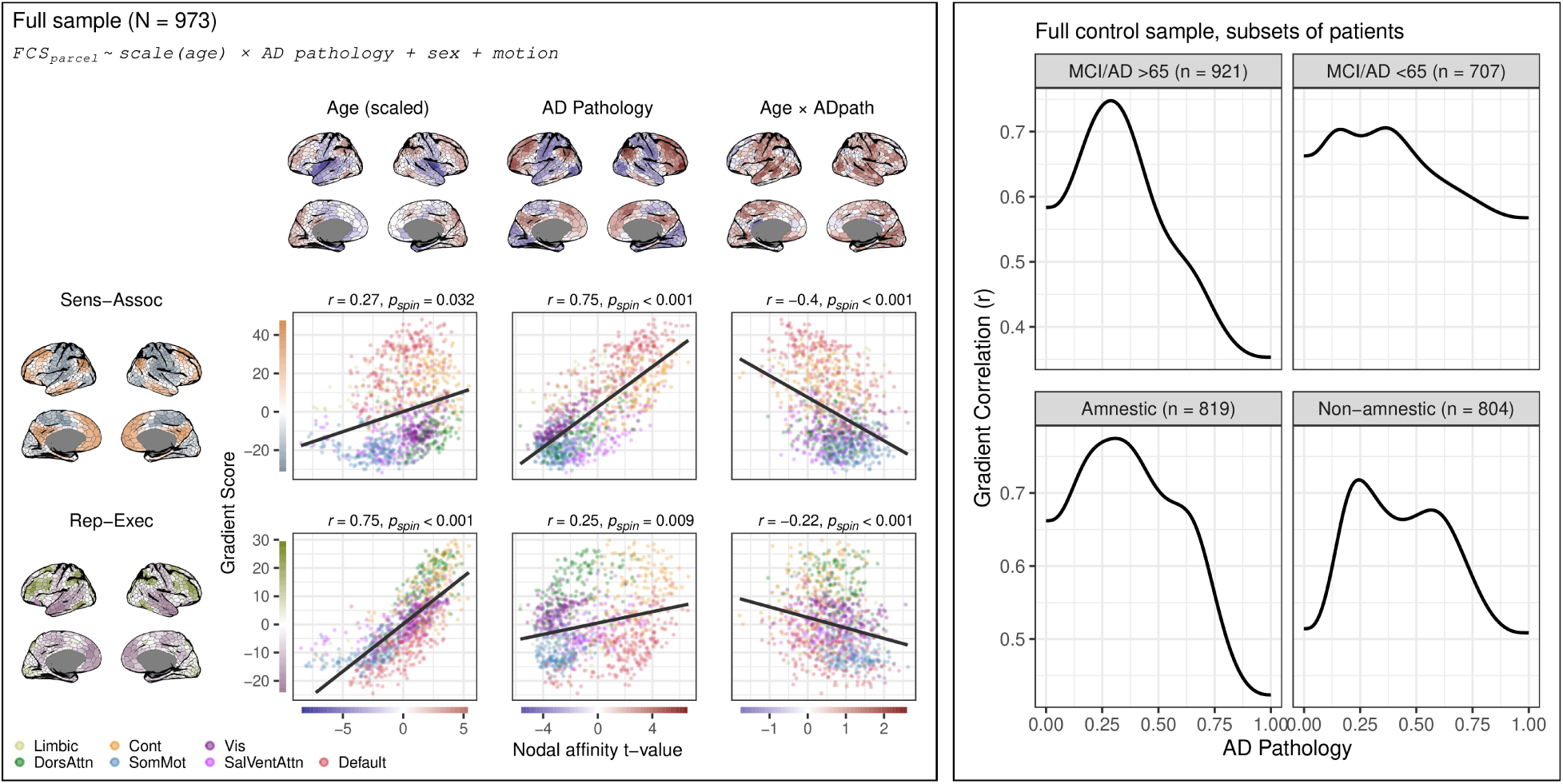
Age attenuates Alzheimer’s disease–related gradient alignment, whereas cognitive phenotype shows limited modulation. Left: Parcel-wise linear regression results in the full sample (N = 973), modeling FCS as a function of age (scaled), AD pathology, and their interaction (controlling for sex and motion). Surface maps show t-values for the main effects and the age × pathology interaction. Scatterplots illustrate the correspondence between parcel-wise t-values and the sensory–association and representational–executive axes, with Pearson correlations and spin-test p-values indicated. The age × pathology interaction shows an inverse alignment with the SA axis, indicating attenuation of pathology-related gradient alignment at older ages. Right: Stratified nonlinear analyses of SA alignment across the AD pathology continuum. Using the full control sample but restricting patient subsets, gradient correlations with the first derivatives of the non-linear parcel-wise slopes were estimated across 100 pathology values from 0 to 1. In the top plot patients were stratified by age and in the bottom plot they were stratified as amnestic/non-amnestic by performance on cognitive domain scores. Domain scores were derived through a bifactor model of cognition visualised in Figure 6. Older patients show a steeper decline in SA axis alignment at higher pathology levels compared to younger patients, whereas both amnestic and non-amnestic groups exhibit broadly similar trajectories. Together, these analyses indicate that age modulates the strength of pathology-related gradient alignment, while broad cognitive phenotype (within typical AD presentation) does not fully account for the observed loss of alignment at later disease stages.

**Figure S3:**
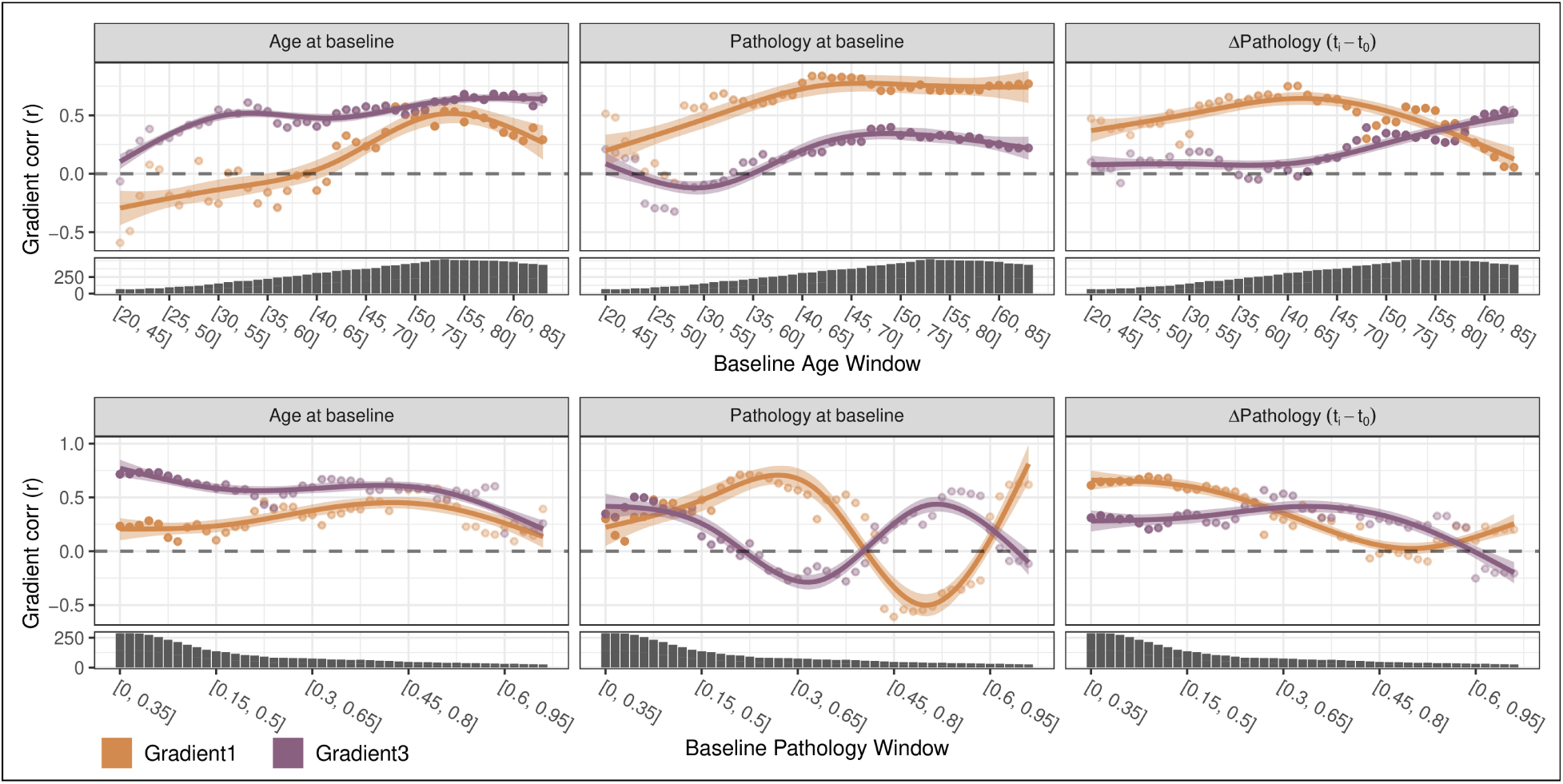
Results of the sliding window analysis from Figure 4 showing the dynamic effect of AD pathology on FC gradient alignment. The correlation values on the scatters are smoothed with generalized additive models. Correlations are shown for each term for both the SA and RE axis. Marginal plots show the sample size for each window. At higher baseline-pathology windows, pathology-related FC effects show a negative association with the sensory–association axis, coinciding with a transient rise in alignment with the representational–executive axis. This crossover indicates that the spatial organization of pathology-related FC changes may shift as pathology advances.

**Figure S4:**
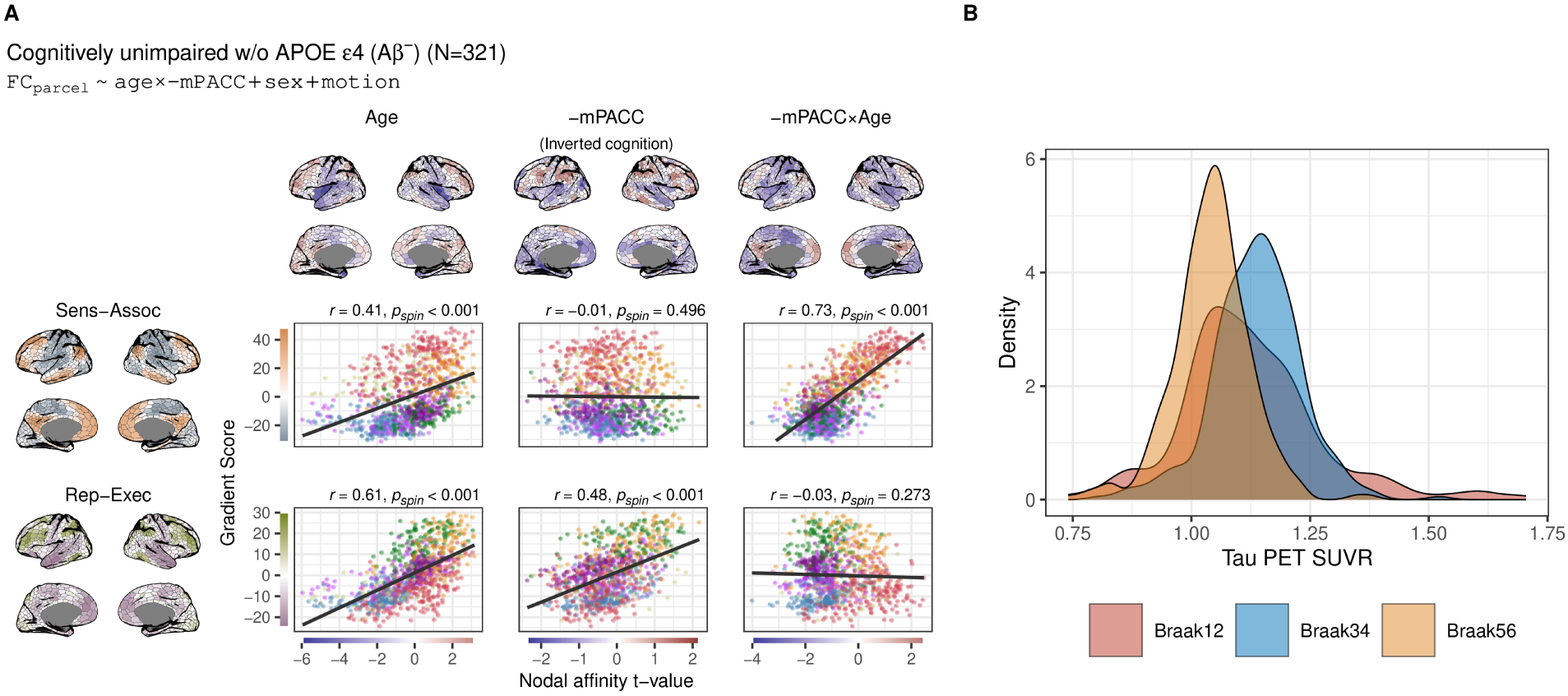
A: Cross-sectional results for cognitively unimpaired Aβ-, *APOE* ε4 non-carriers. The model includes age, inverted cognition (-mPACC), and an age × cognition interaction (with centered predictors and inverted cognition scores for interpretability). Unlike the main analysis presented in Figure 5, this analysis does not adjust for pathology. The age effects observed here partially reflect the effects previously attributed to pathology, revealing a relationship with the SA axis that diminishes when adjusting for pathology. Cortical maps display t-values from nodal linear regression models, while scatter plots show the relationships between the t-values and gradient scores, colored by network membership. The relationship was quantified using Pearson correlation and significance assessed using a spin test (one-sided, uncorrected; see Methods). B: Distribution of tau PET SUVR from Braak I-II, Braak III-IV and Braak V-VI in cognitively unimpaired Aβ-, *APOE* ε4 non-carriers.

**Figure S5:**
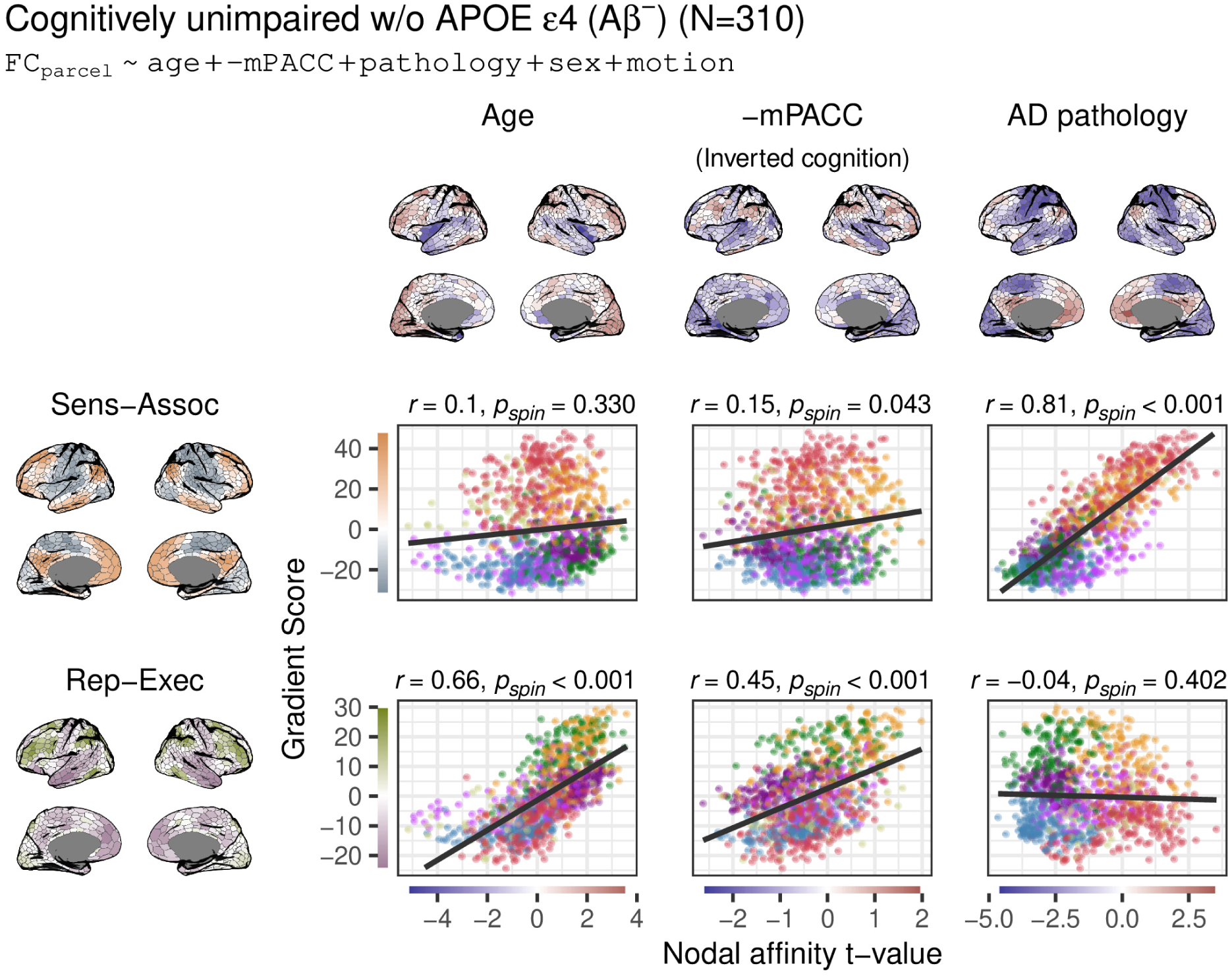
Cross-sectional results for cognitively unimpaired Aβ-, *APOE* ε4 non-carriers. This model is without interaction between cognition and age.

**Figure S6:**
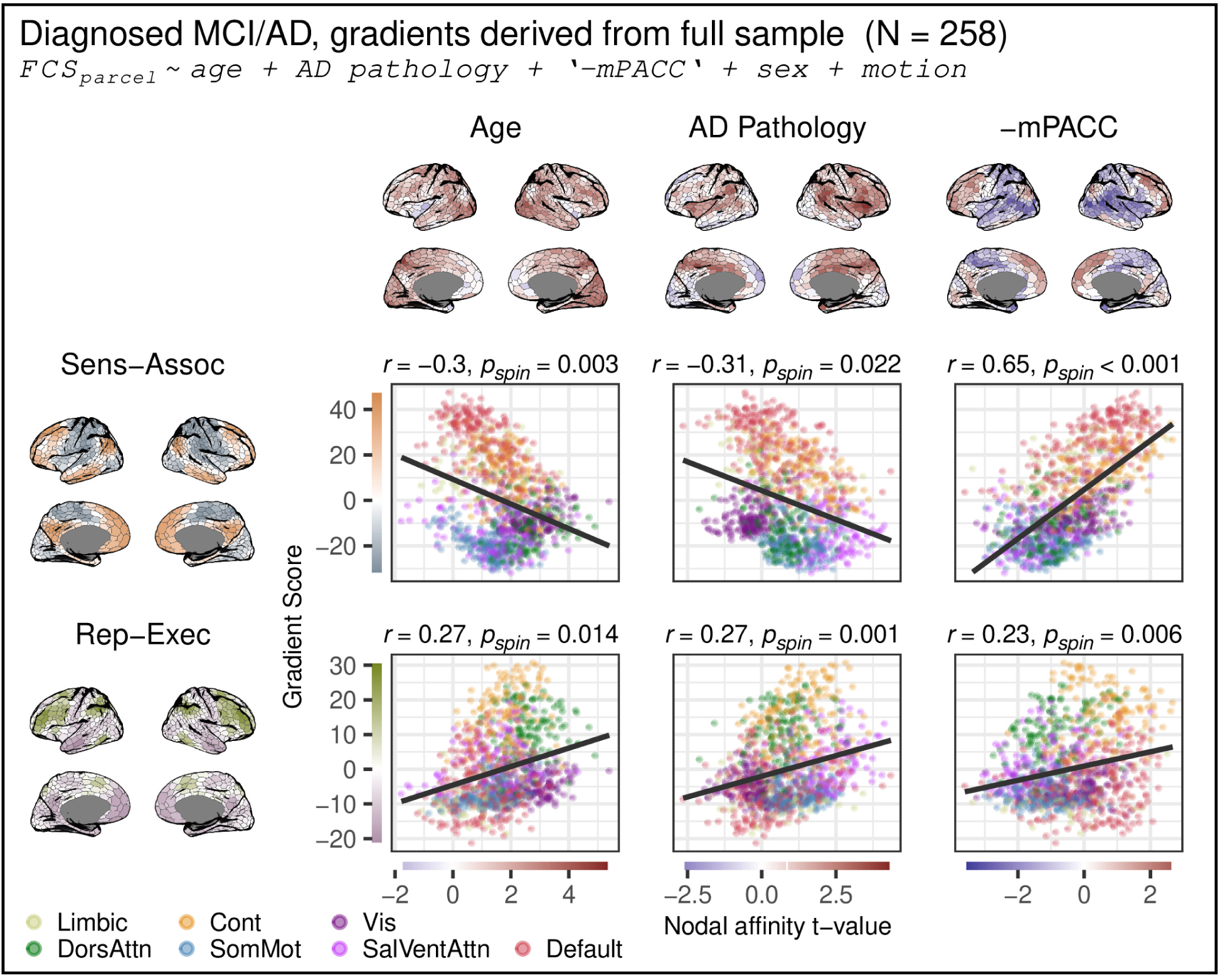
To investigate whether the lost association between AD pathology and the sensory association axis in the cognitively impaired subgroup analysis was related to a bias in deriving the gradients only from cognitively healthy subjects below the age of 60, we derived gradients from the full sample and ran the same analysis. This produced consistent results.

**Figure S7:**
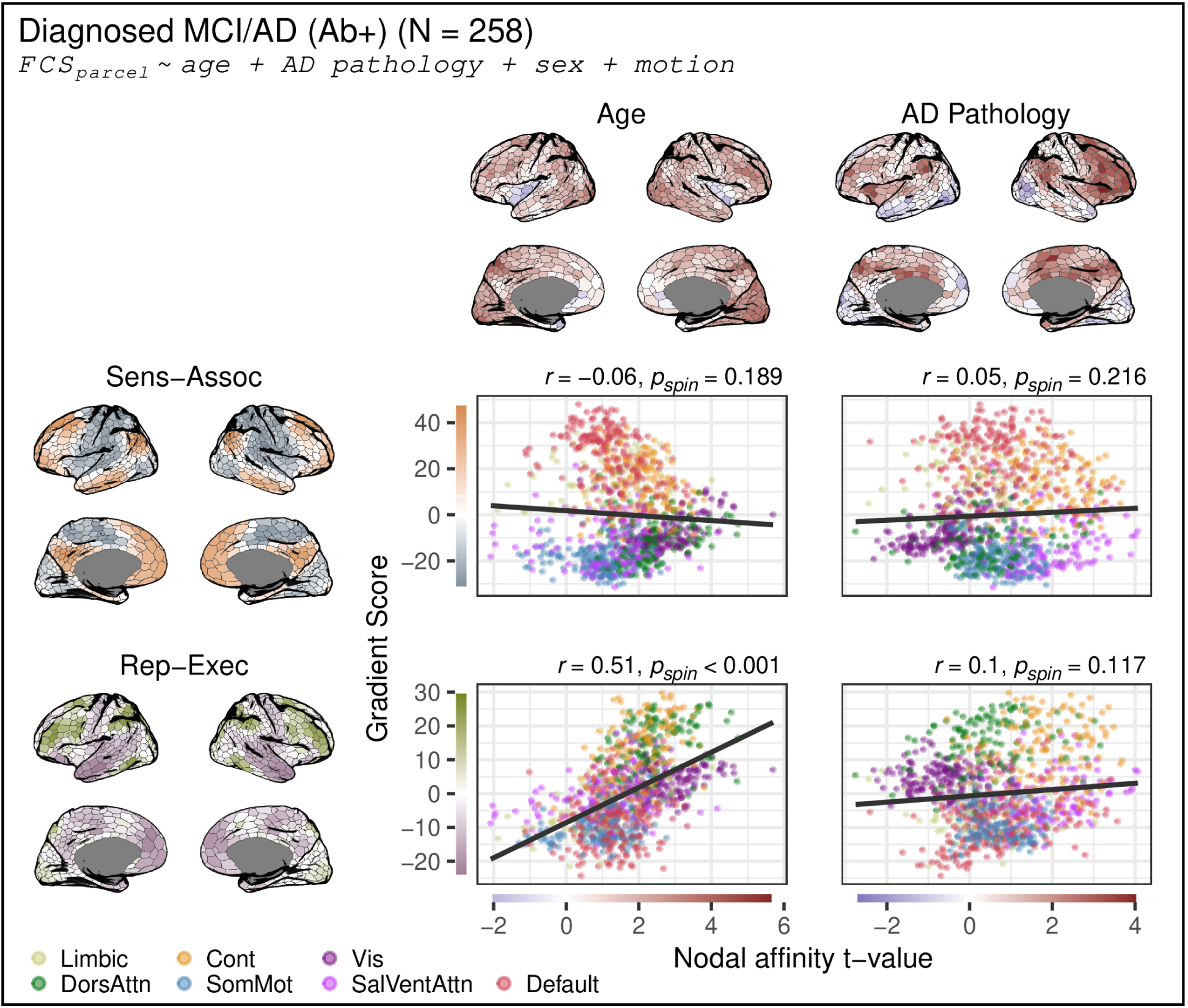
Cross-sectional results within the MCI and AD group without cognition. This analysis revealed that AD pathology showed no relationship to the SA axis even when not accounting for cognition, within this group.

**Figure S8:**
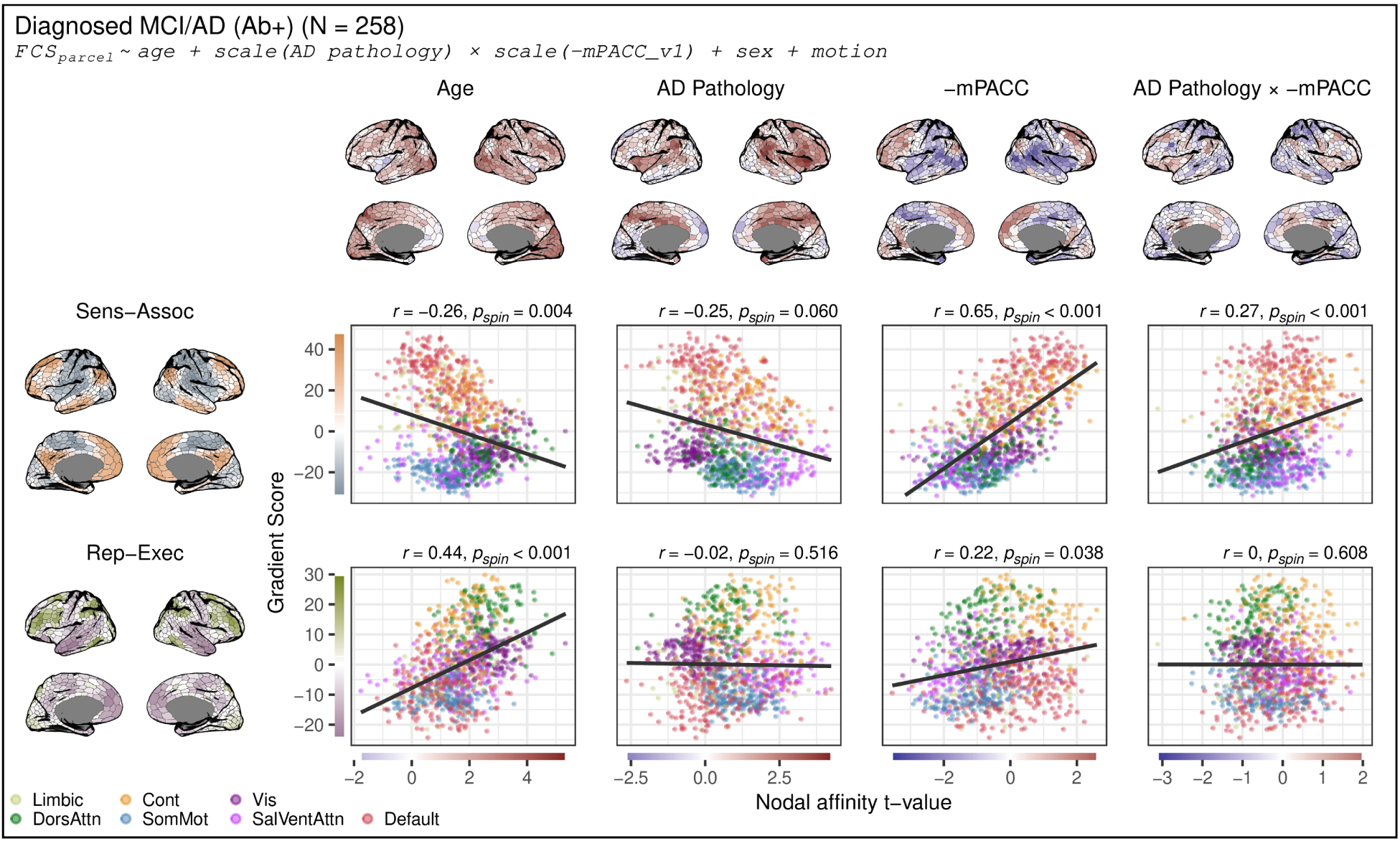
Cross-sectional analyses within patients with mild cognitive impairment (MCI) and Alzheimer’s disease (AD), interacting AD pathology and cognition revealed that this did not affect the main effect of cognition. Predictors have been centred and cognition scores inverted for interpretability. Cortical maps display t-values from nodal linear regression models, while scatter plots show the relationships between the t-values and gradient scores, colored by network membership. The relationship was quantified using Pearson correlation and significance assessed using a spin test (one-sided, uncorrected; see Methods).

**Figure S9:**
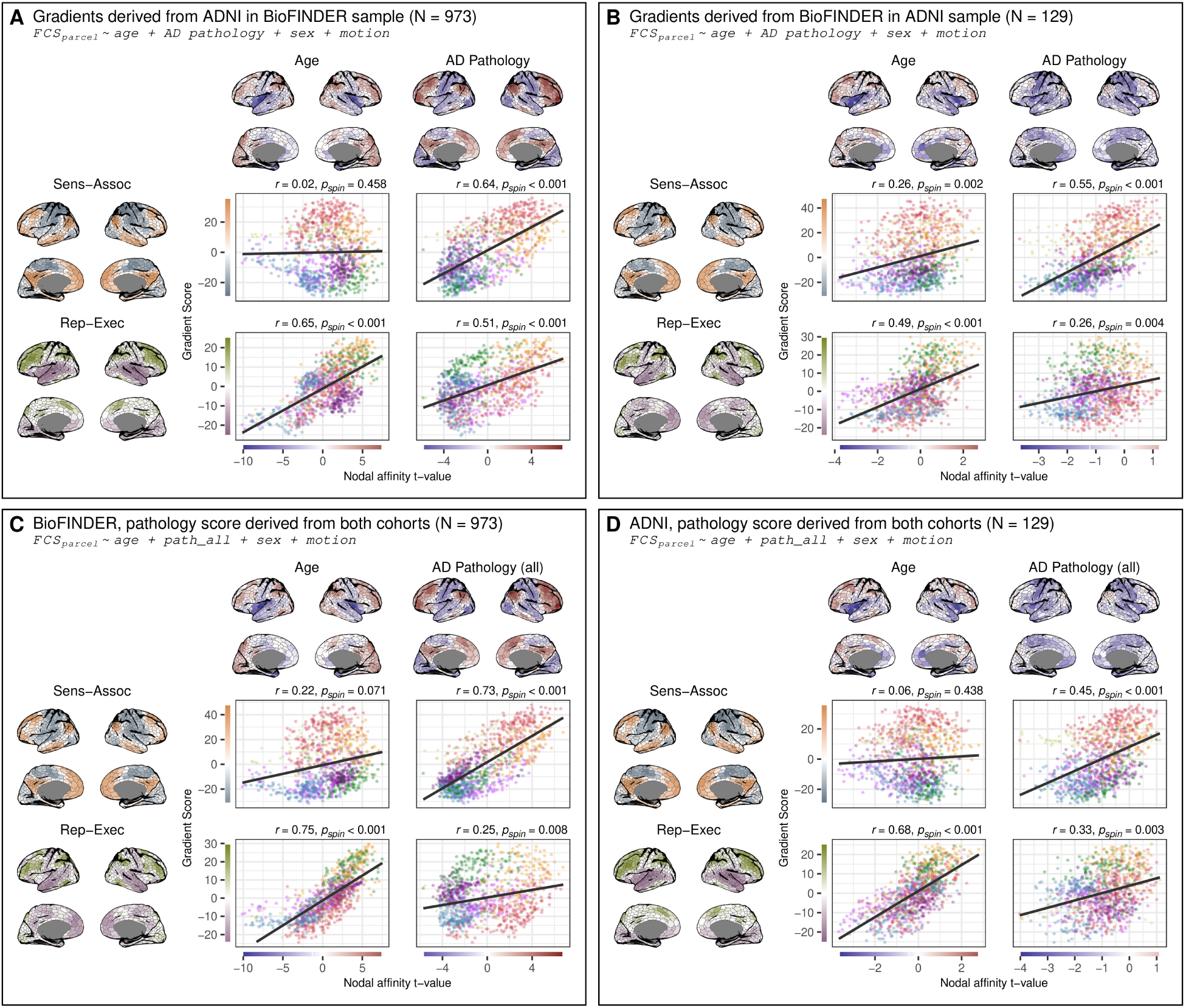
Main results but using gradients derived from ADNI to interpret the t-maps in BioFINDER (A) and the gradients derived from BioFINDER to interpret the t-maps from ADNI (B). To investigate whether the calculation pathology scores within each cohort affected the results we derived a pathology score after pooling data from the two samples and running the main analysis again in both BioFINDER (C) and ADNI (D). Results were comparable with the originally derived pathology score within each cohort.

**Figure S10:**
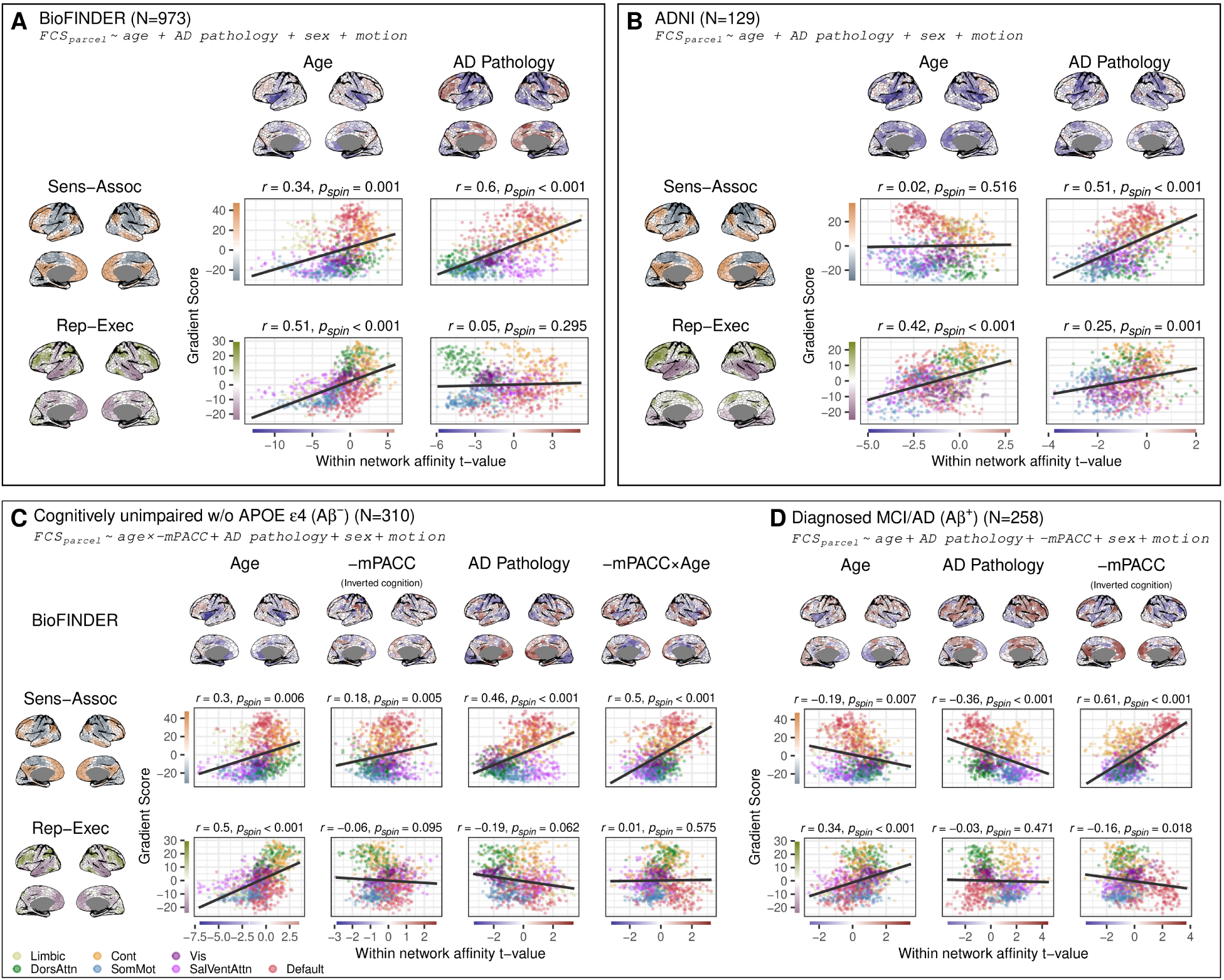
Analyses replicated with within-network affinity. Within-network affinity was calculated as the average affinity between parcels within the same Yeo network^38^. Results are shown for BioFINDER (A), ADNI (B). How cognitive status seems to modify these relationships are shown in (C) and (D). Cortical maps display t-values from nodal linear models, while scatter plots show the relationships between t-values and gradient scores, colored by network membership. The relationship was quantified using Pearson correlation and significance assessed using a spin test (one-sided, uncorrected; see Methods).

**Figure S11:**
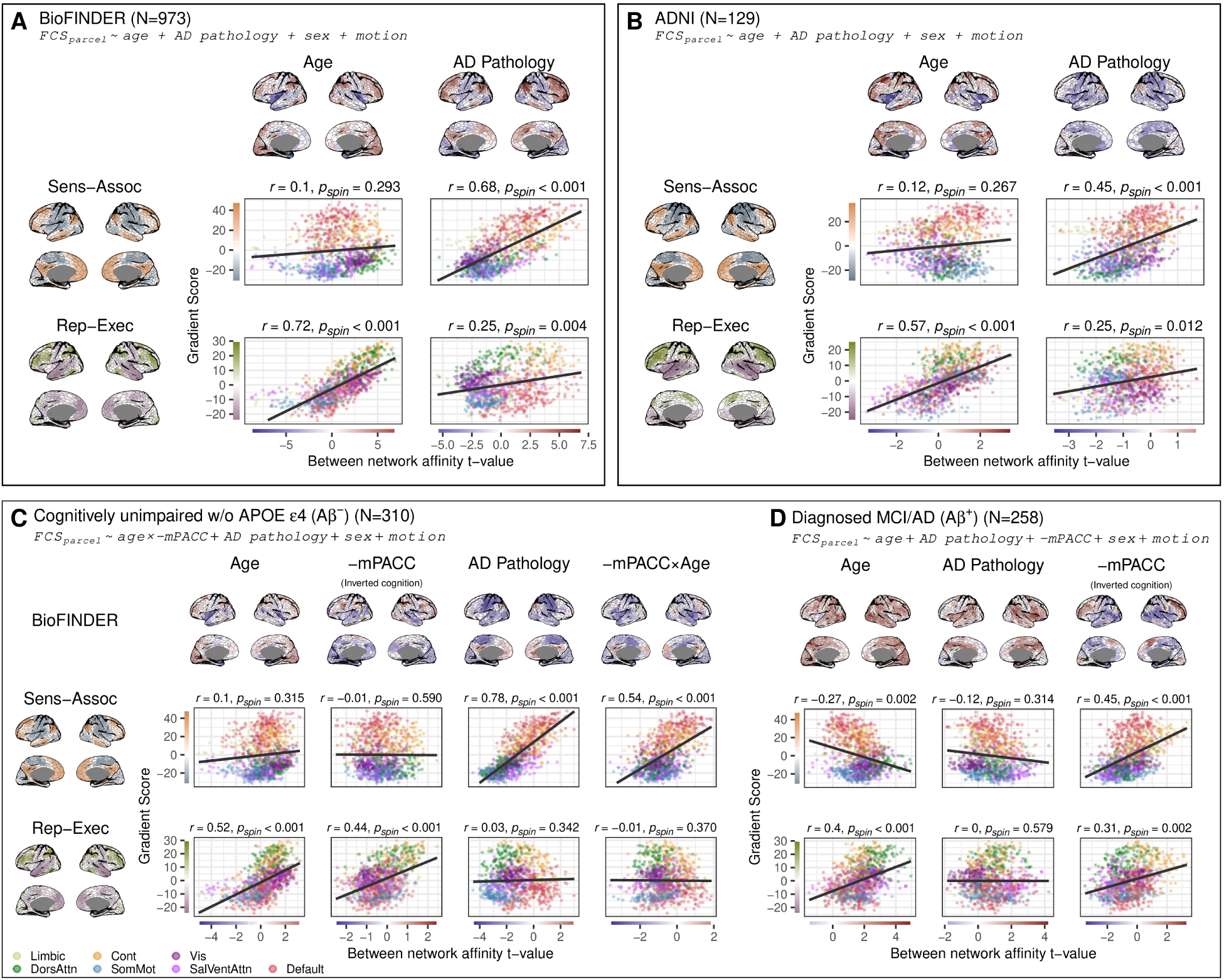
Analyses replicated with between-network affinity. Between-network affinity was calculated as the average affinity to parcels outside of a parcel’s own Yeo 7 network^38^. Results are shown for BioFINDER (A), ADNI (B). How cognitive status seems to modify these relationships are shown in (C) and (D). Cortical maps display t-values from nodal linear models, while scatter plots show the relationships between t-values and gradient scores, colored by network membership. The relationship was quantified using Pearson correlation and significance assessed using a spin test (one-sided, uncorrected; see Methods).

**Figure S12:**
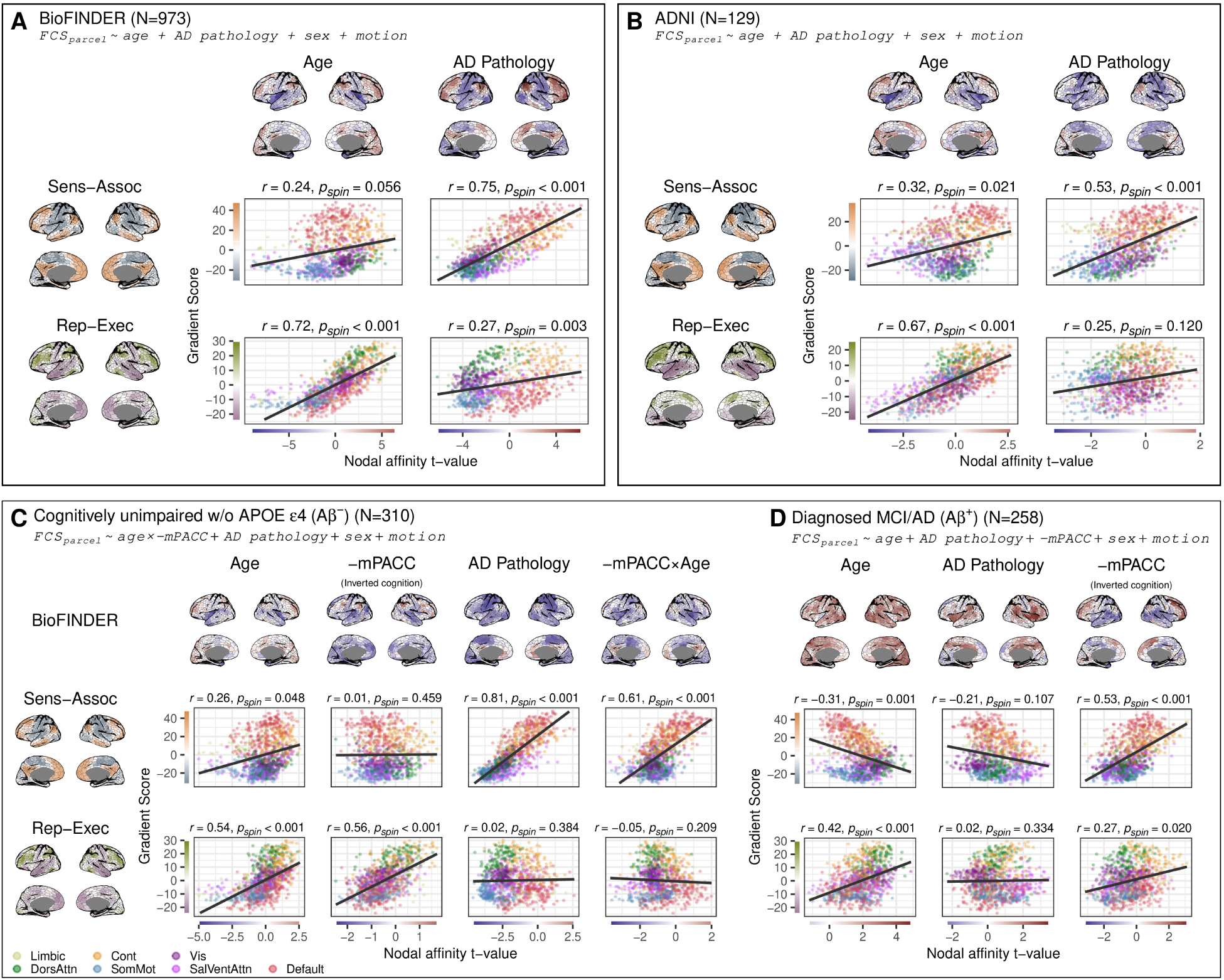
Analyses replicated with nodal affinity calculated using Pearson correlation instead of cosine similarity and no thresholding demonstrate consistent results with the main analyses (Figure 2 and Figure 5). Results are shown for BioFINDER (A), ADNI (B). How cognitive status seems to modify these relationships are shown in (C) and (D). Cortical maps display t-values from nodal linear models, while scatter plots show the relationships between t-values and gradient scores, colored by network membership. The relationship was quantified using Pearson correlation and significance assessed using a spin test (one-sided, uncorrected; see Methods).

